# A Fast Lasso-Based Method for Inferring Pairwise Interactions

**DOI:** 10.1101/2021.01.28.428698

**Authors:** Kieran Elmes, Astra Heywood, Zhiyi Huang, Alex Gavryushkin

## Abstract

Large-scale genotype-phenotype screens provide a wealth of data for identifying molecular alternations associated with a phenotype. Epistatic effects play an important role in such association studies. For example, siRNA perturbation screens can be used to identify pairwise gene-silencing effects. In bacteria, epistasis has practical consequences in determining antimicrobial resistance as the genetic background of a strain plays an important role in determining resistance. Existing computational tools which account for epistasis do not scale to human exome-wide screens and struggle with genetically diverse bacterial species such as *Pseudomonas aeruginosa*. Combining earlier work in interaction detection with recent advances in integer compression, we present a method for epistatic interaction detection on sparse (human) exome-scale data, and an R implementation in the package `Pint`. Our method takes advantage of sparsity in the input data and recent progress in integer compression to perform lasso-penalised linear regression on all pairwise combinations of the input, estimating up to 200 million potential effects, including epistatic interactions. Hence the human exome is within the reach of our method, assuming one parameter per gene and one parameter per epistatic effect for every pair of genes. We demonstrate `Pint` on both simulated and real data sets, including antibiotic resistance testing and siRNA perturbation screens.

## 1. Introduction

Epistatic gene interactions have practical implications for personalised medicine, and synthetic lethal interactions in particular can be used in cancer treatment [3]. Discovering these interactions is currently challenging [23, 17, 10, 12], however. In particular, there are no methods able to automatically infer interactions from genotype-phenotype data at the human genome scale.

For a given number of genes there are exponentially many potential interactions, complicating computational methods. If we restrict our attention to pairwise effects, it is possible to experimentally knock out particular combinations of genes to determine their combined effect [9]. This approach does not scale to the approximately 200 million pairwise combinations possible among human protein coding genes, however. We instead consider inferring pairwise interactions from large-scale genotype-phenotype data. These include mass knockdown screens, in which we suppress a large number of genes simultaneously, and attempt to measure the resulting phenotypic effect.

We have shown in [12] that a lasso-based approach to inferring interactions from an siRNA perturbation matrix is a feasible method for large-scale interaction detection. In this additive model, we assume fitness is a linear combination of the effects of each gene’s effect, and the effect of every combination of these genes. For the sake of scalability, we consider only individual and pairwise effects, and assume gene suppression is strictly binary. The fitness difference *f* (compared to no knockdowns) in an experiment e is then the sum of individual and pairwise effects 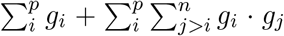, where *g_i_* = 1 if gene *i* is knocked down, 0 otherwise. With sufficiently many such mass-knockdowns, we can infer pairwise interactions by finding the pairs of genes whose effect is not the sum of the effects of each gene individually.

Neither of the previously tested inference methods for this model, `glinternet` and `xyz`, are effective at the genome-scale however. `glinternet` suffers from prohibitively long running times,^1^ and `xyz` does not accurately predict effects in our larger simulations. Our aim is to fit a model including all *p* ≈ 20, 000 human protein-coding genes, with as many as *n* = 200, 000 siRNAs. Doing so requires the development of new methods and software.

We have developed an R-package that is able to perform lasso regression on all pairwise interactions on the same one thousand gene screen in twenty seconds, and is able to fit a genome-scale data set with 19, 000 genes and 67, 000 siRNAs in under two hours using a single eight-core CPU. This is made possible by taking into account that our input matrix *X* is both sparse and strictly binary. Our package, `Pint`, is available at github.com/biods/pint.

To perform lasso-based regression on this matrix, we begin with an existing fast algorithm, parallelise it, and adapt it for use on our binary perturbation matrices. We provide a detailed explanation of this implementation, followed by the scalability analysis, below. We also perform a simulation study to compare our method’s scalability with known methods, and analyse two large-scale experimental data sets.

In the first, an siRNA perturbation screen from [31], we search for pairs of genes that have an epistatic effect when simultaneously silenced. Out of five top interactions identified by our method, two are known protein interactions and three appear to be novel.

The second data set is composed of genetic variants identified in the intrinsically antibiotic resistant bacteria *Pseudomonas aeruginosa. P. aeruginosa* is an opportunistic pathogen found in a variety of environments and is a leading cause of morbidity and mortality in immunocompromised individuals or those with cystic fibrosis [16, 24]. *P. aeruginosa* is known to acquire adaptive antibiotic resistance in response to long term usage of antibiotics associated with chronic infections [5, 27, 28]. The genomes included in that data set are from strains that have been isolated from chronic and acute infections as well as environmental samples. The minimum inhibitory concentration for the antibiotic Ciprofloxacin has been used as the phenotypic marker for this dataset. Ciprofloxacin belongs to the fluoroquinolone class of bacteriocidal antibiotics that targets DNA replication and is one of the most widely used antibiotics against *P. aeruginosa*[34]. Our findings identified 16 pairs of interactions, most of which were found in genes that are important in biofilm formation and maintenance, a characteristic of intrinsically antibiotic resistant bacteria.

## 2. Methods

Our goal is to estimate both the main effects *β*_1_,…,*β*_p_, and the interaction effects *β*_1,2_,…, *β*_*P*-1,*P*_ where pairs of genotypes are simultaneously perturbed. As an example, consider pairwise effects in a siRNA perturbation screen. We can estimate the effect of both silencing individual genes (*β*_1_,…) and pairs of genes simultaneously (*β*_1,2_,…). To do this we add a column for each pair of genes, converting the siRNA matrix **X** ∈ {0,1}^n×p^ into the pairwise matrix **X**_2_ ∈ {0,1}^n×p’^, where 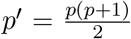. This model includes all pairwise interactions and fitting it is equivalent to finding epistasis as in [12]. The same construction applies to any binary genotype-phenotype data, and the effect *β_a,b_* will always estimate the simultaneous effect of both genes *a* and *b*.

We construct the matrix **X**_2_ as follows. For every column *i* from 0 to *n* we take every further column *j* from *i* + 1 to *n* and form a new column by taking the bit-wise and over all elements of the columns *i* and *j* (Fig. 1).

**Figure 1.**
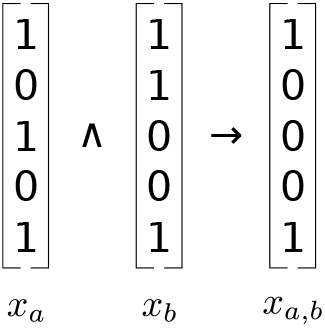
Creation of pairwise siRNA effect columns

This gives us the complete pairwise matrix **X**_2_, shown in Fig. 2.

**Figure 2.**
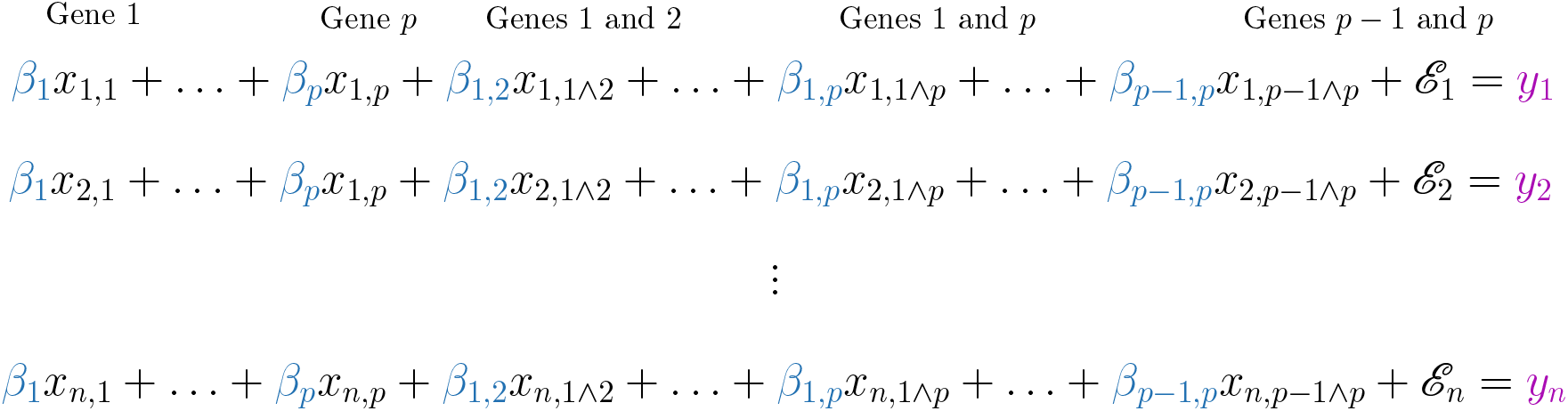
Matrix of Pairwise siRNA effects

### 2.1. Cyclic Linear Regression

Our approach to lasso regression is based on a cyclic coordinate descent algorithm from [15], as described in [40]. This method begins with *β_j_* = 0 for all *j* and updates the beta values sequentially, with each update attempting to minimise the current total error. Here this total error is the difference between the effects we have estimated and the fitness we observe, given the genes that have been knocked down. Where *y_i_* is the ith element of **Y**, *β_j_* is the *j*th element of *β*, and *X_ij_* is the entry in the matrix **X**_2_ at column *j* of row *i*, the error is the following.

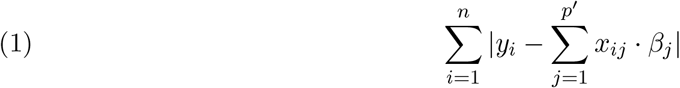

The error affected by a single beta value (Eq. (5)) can then be minimised by updating *β_k_* with the following:

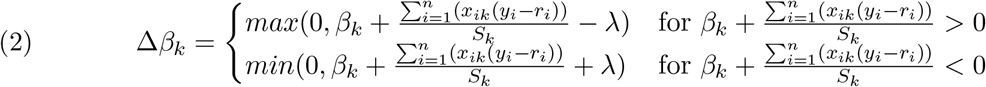

We cyclically update each *β_k_* until the solution converges for a particular lambda, reduce the value of lambda, and repeat. See Appendix A for the full derivation and algorithm. Storing the matrix in a sparse column format, this implementation scales up to *p* = 1,000. It would still take several days and use terabytes of memory for *p* = 20,000. To overcome this, we compress the matrix, and parallelise the beta updates (Section 2.3 and Appendix D).

### 2.2. Choosing Lambda

The lasso penalty requires a regularisation parameter lambda. This parameter determines the extent to which we penalise large beta values, and can range from allowing all values (λ = 0) to allowing only zero (λ → ∞). Choosing the correct value of lambda is essential if we want to include only the significant effects. This is typically done by choosing an initial value sufficiently large that all beta values will be zero and gradually reducing lambda, fitting the model for each value until a stopping point chosen with K-fold cross-validation [14]. Cross-validation requires fitting each lambda value *K* times, however, significantly increasing the runtime. We instead provide two options for choosing lambda in our package. First, we can choose lambda such that the number of non-zero effects is small enough for OLS regression. In our package, this is called Limited-*β* and a default limit is 2,000. Alternatively, we implement a fast method for empirically choosing a reasonable stopping point, the adaptive calibration lambda selection method from [8]. Both of these methods are significantly faster than crossvalidation, although using adaptive calibration we tend to predict very few non-zero effects. The best empirical results are generally achieved with the Limited-*β* approach, and we use this for the remainder of the paper. A detailed explanation of each and their performance impact can be found in Appendix C.

### 2.3. Compression

To reduce memory usage and the time taken to read each column with larger input data, we compress the columns of **X**_2_. Because we read the columns sequentially, we replace each entry with the offset from the previous entry. This reduces the average entry to a relatively small number, rather than the mean of the entire row. These small integers can then be efficiently compressed with any of a range of integer compression techniques (Fig. 3), a subject that has been heavily developed for Information Retrieval. We compare a number of such methods, including the Simple-8b algorithm from [35] (which we implement and use in our package) in Appendix B.

**Figure 3.**
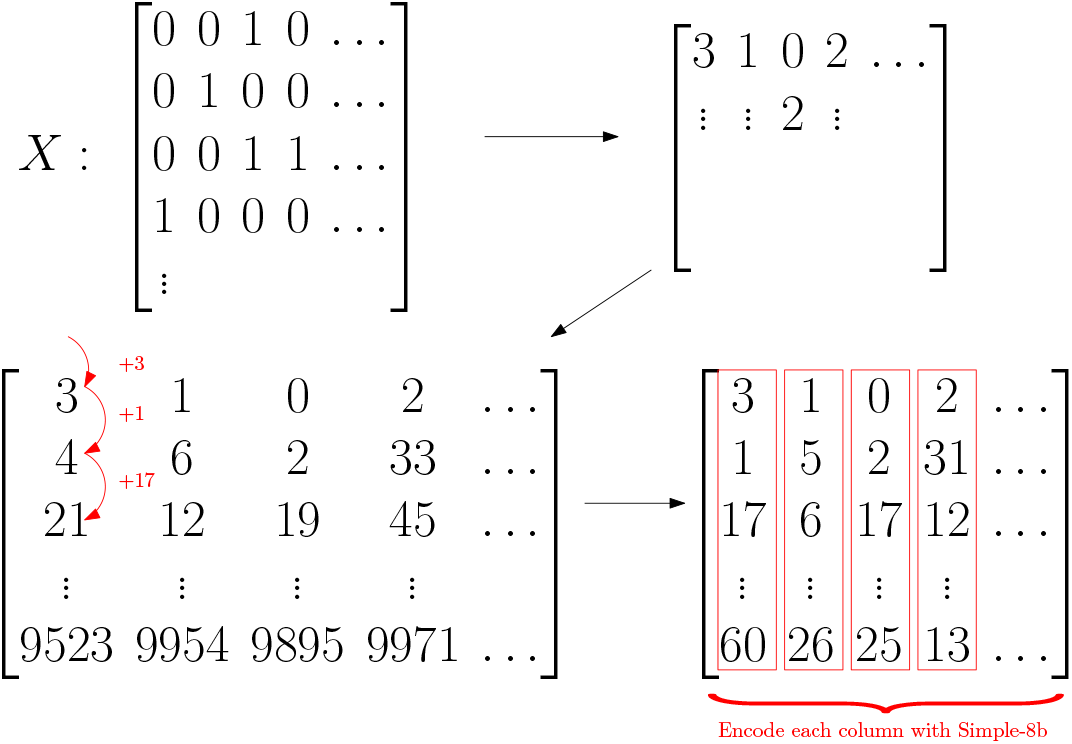
Compression of the sparse **X**_2_ matrix.

### 2.4. Parallelisation

While it is trivial to parallelise the update of a single *β* value, doing so does not improve performance in practice, due to poor cache usage (see Appendix D for details). We instead parallelise our method by assigning whole columns of the compressed **X**_2_ matrix to threads in sections. Each thread is responsible for updating the *β* values corresponding to its columns, and is therefore the only thread reading it’s section of the **X**_2_ matrix.

Simultaneously updating columns with entries in the same row leads to over-compensating for these entries. This can harm performance or in the worst case prevent convergence entirely (Appendices D.2, D.5 and D.5.1). To avoid this, it suffices to ensure that threads do not frequently update the same columns at the same time. We achieve this by shuffling the order each thread updates it’s columns every iteration. While it is in principle still possible to update enough overlapping threads in parallel to cause problems, Bradley et al. [6] show that this is rarely a problem in practice.

Updates to the shared *β* values are atomic, and every thread needs read access to all *β* values. This limits our method to use on shared memory systems and results in poor performance on NUMA systems. In practice we can only effectively use a single CPU socket, and we have tested this up to eight cores. Fig. 4 summarises the shuffled parallel implementation and it’s scalability. For the full details of the parallel implementation see Appendix D.

**Figure 4.**
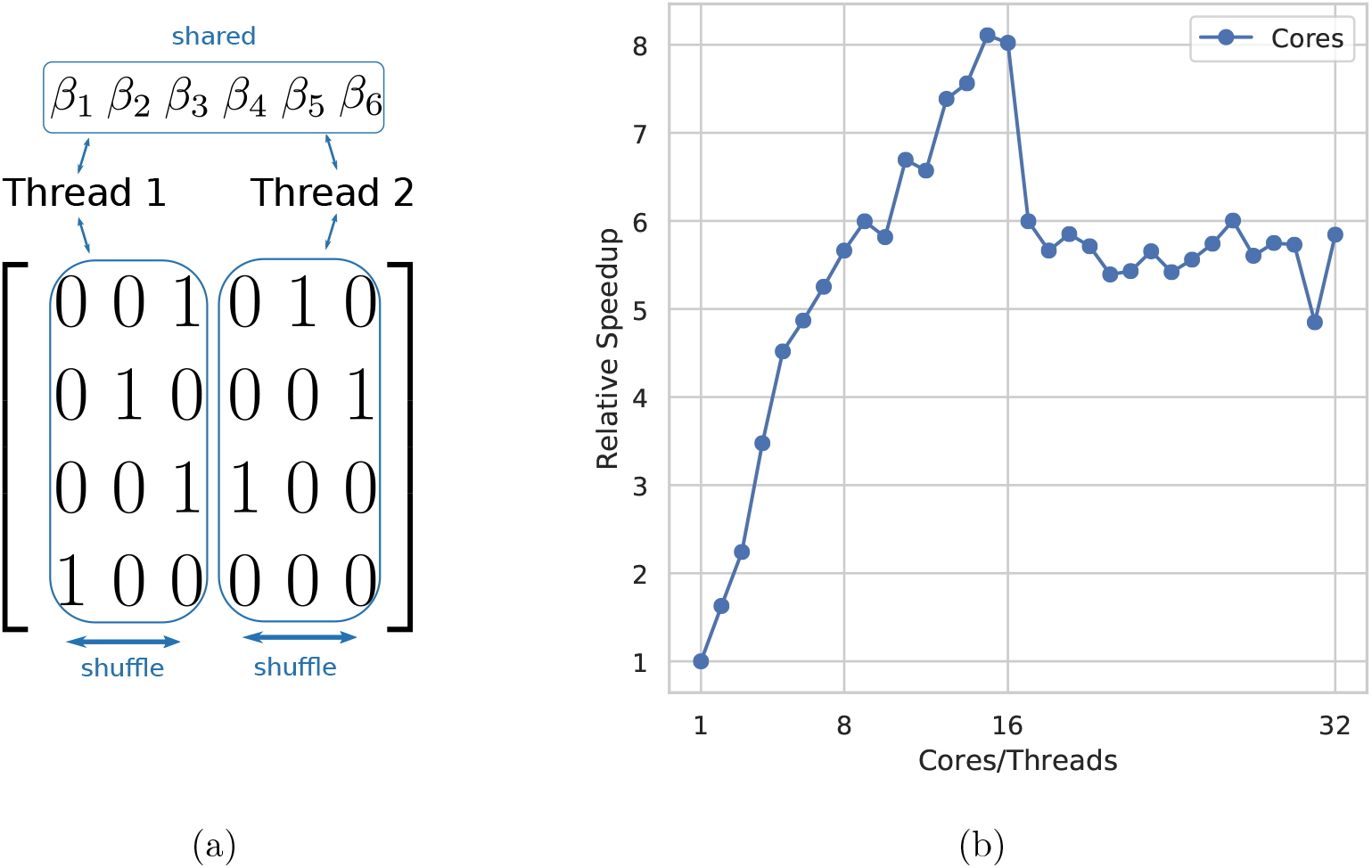
(a) Each thread is assigned a set of columns, which is then shuffled every iteration. (b) Relative speedup as the number of cores used increases, running on a dual 8 core/16 thread NUMA system. Cores 1-8 are separate cores on node 1, 8-16 are SMT threads on the same cores. Cores 17-24 are separate cores on the second NUMA node, and 15-32 are SMT threads on those cores.

### 2.5. Limited Interaction Neighbourhoods

When searching for interactions within a large sequence, it may be acceptable to limit the search to pairs that are relatively close on the genome. In a study of epistatic interactions in yeast by Puchta et al. [30] the strength of negative interactions decreases as distance between gene positions on the sequence increases. The median distance between pairs in the hundred strongest interactions was only eighteen nucleotides.

Limiting interactions to those within some distance d drastically reduces both the time and space requirements. Instead of Θ(*p*^2^n), the size of the interaction matrix becomes Θ(pdn). Similarly, an iteration of Algorithm 3 would require only Θ(pdn) operations. For d ≪ *p* this is a significant reduction. Limiting the interaction search distance to 100 positions, we could process a set of 30,000 genes and 200,000 siRNAs using approximately 16*GB* of memory, assuming a comparable density of interactions to our testing data. Such a search could be performed directly on a laptop, without requiring access to a large server. The biological implications of this restriction should be carefully considered before its use, however.

### 2.6. Data

We prepared two experimental data sets to evaluate our method and test the scalability of our implementation. The first is an siRNA perturbation screen in which siRNAs targeting kinases are applied to an infected human cell line. We predict off-target effects across the entire exome, and use this larger set for our analysis.

The second data set contains single nucleotide variants (SNVs) from 259 isolates of *Pseudomonas aeruginosa*, and associated minimum inhibitory concentration (MIC) of Ciprofloxacin.

#### 2.6.1. InfectX siRNA Data

To demonstrate our method on real genome-scale data, we use the vaccinia group from InfectX [31]. This set contains 204,288 siRNA perturbations in the presence of the vaccinia virus. This set is significantly larger than the mock group (siRNA perturbations with no pathogen present). Off-target effects are prediction using Risearch2 [1]. We include a gene as an off-target effect whenever there is a match between the siRNA seed region and some component of an mRNA for that gene (taken from [18]). We use an energy cutoff of −20 and match the entire siRNA, not only the 3’ UTR, as suggested in [1].

We then form a matrix of off-target effects with columns for each gene, and rows for each siRNA as in [12]. An entry *i, j* in this matrix is one if and only the predicted effect of siRNA *i* on gene *j* is greater than zero. All other entries are zero. Our fitness vector **Y** is the result of B-scoring then Z-scoring the number of cells in the well, to remove systematic within-plate effects and experimentally introduced cross-plate biases. B-scoring corrects for biases across the entire plate, and Z-scoring then normalises each well’s score with respect to the rest of it’s plate.

#### 2.6.2. Antibacterial Resistance

SNVs from 259 isolates of *Pseudomonas aeruginosawere* sequenced using illumina technologies (IPCD isolates on MiSeq and QIMR isolates on HISeq). SNV’s from raw reads were mapped to the reference genome PAO1 using Bowtie2 (v. 2.3.4) [21] read aligners. Variant reports were then read into a python script which sorted the reports into a table. The table was set up so that each isolate was represented as a row and the presence / absence of each SNV was along the columns. Only genomes that had associated MIC values were included. The resulting table contains 259 rows and over 700, 000 columns.

Since our method considers *p*^2^ interactions, the scale of this data presents a problem. Including all > 700,000 columns, we would need to store over 250 billion interaction columns, each with up to 259 entries. Even if every column fits into a single 64-bit word, simply storing the compressed matrix would require on the order of two terabytes of memory. We instead reduce this to a more manageable scale, by removing all duplicated columns, and then any of the remaining columns that have less than 30 entries. Note that this is likely to remove point mutations occurring from acquired resistance, and effects that are always found in the same isolates cannot be distinguished. While it may be possible to address these limitations we do not attempt to do so here. There are simply too many interactions (over 200 billion) among the full set of variants for our current implementation. After these reductions we have a more tractable 259 × 75, 715 entry matrix, sufficiently small that all approx. 5.7 billion effects and interactions can be processed using under 250GB of memory.

#### 2.6.3. Data Sources

*P. aeruginosagenome* sequences were selected from strains whose MIC values (Ciprofloxacin) were known. 167 genomes were sourced from the publicly available IPCD International Pseudomonas Consortium Database [19] and 92 genomes were from QIMR Brisbane Australia [20]. The IPCD data consisted of 2 x 300 bp MiSeq reads whilst the QIMR data was 2 x 150 bp reads. The MIC values were obtained as a combination of e-test strips [32] and plate-based assays [33].

## 3. Results

In this section, we summarise the results of a simulation study we carried out to compare our method against existing approaches. We also demonstrate our method on two large-scale experimental data sets. Note that both sets are too large to attempt using known approaches such as `glinternet` for comparison. In both cases the true interactions are unknown, making the true accuracy of our method in these cases difficult to determine. We nonetheless include these as reasonable examples of cases in which our method is applicable and validate the results by comparing them with known protein interactions [36].

### 3.1. Simulation Performance

Our method aims to have comparable precision and recall to the best performing approach in our previous work [12] while scaling to much larger data sets. To evaluate the accuracy of our method, we compare precision and recall of our method with `glinternet`, the most accurate of the methods tested [12].

Since we achieved the best results only using `glinternet` for variable-selection, then fitting the non-zero beta values with ordinary least squares (OLS) regression, we do the same here. We use Pint in the same way and restrict to the first 2, 000 non-zero beta values, rather than using adaptive calibration, which returns too few columns for the OLS regression step.

**Table 1.** Runtime comparison between our method and `glinternet`.

| **X** matrix size | **glinternet** time (s) | **Pint** time (s) |
| --- | --- | --- |
| $n = 100, p = 1,000$ | 178 | 2.00 |
| $n = 1,000, p = 10,000$ | 4807 | 27.6 |

**Table 2.** Infectx proposed interactions

| Gene Names |  | Estimated Effect | p-value |
| --- | --- | --- | --- |
| TTN | KMT2D | 0.085 | 4.35e-05 |
| TTN | PLEC | 0.053 | 1.95e-2 |
| TTN | TTC7B | 0.068 | 2.01e-3 |
| TTN | OBSCN | 0.137 | 8.00e-11 |
| TTN | CDH23 | -0.022 | 1.00e-1 |

Testing with the same data as in [12], our method is able to identify significantly more correct interactions than `glinternet` (Fig. 5). Precision is largely comparable, with a few outliers in which we see significantly more false positives with our method (Fig. 5a). The run time is orders of magnitude faster than `glinternet`, typically taking 20 to 30 seconds rather than several hours (Table 1 and Fig. 5c). To test the scalability of our implementation, we also run it with the same 2, 000 effect limit on a much larger data set. With p ≈ 27,000, n ≈ 30,000, using 16 SMT threads on a single eight core CPU, we propose 97 main effects and 236 interactions in one and a half hours.

**Figure 5.**
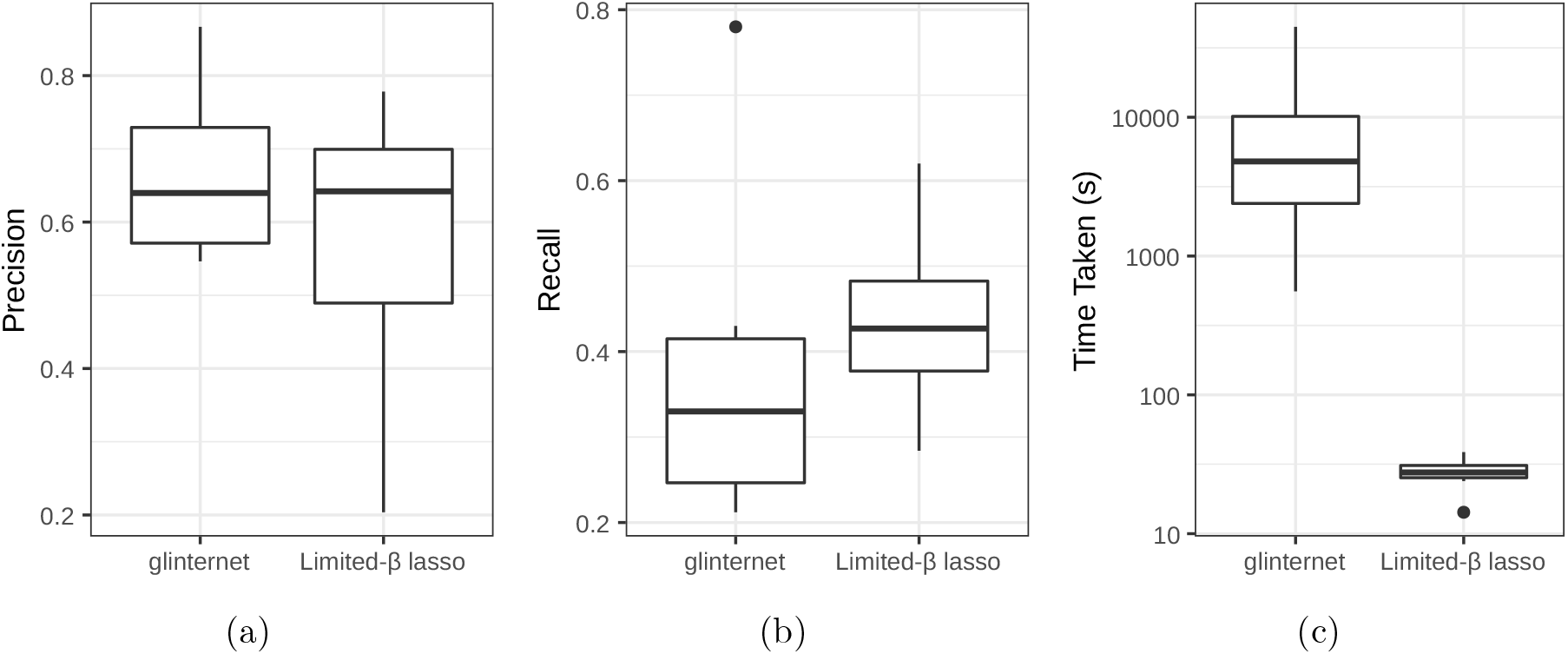
Searching for interactions with glinternet vs. our shuffled compressed lasso, using *p* = 1, 000, *n* = 10,000 data from [12]. (a) Precision. (b) Recall. (c) Time taken (log scale).

**Figure 6.**
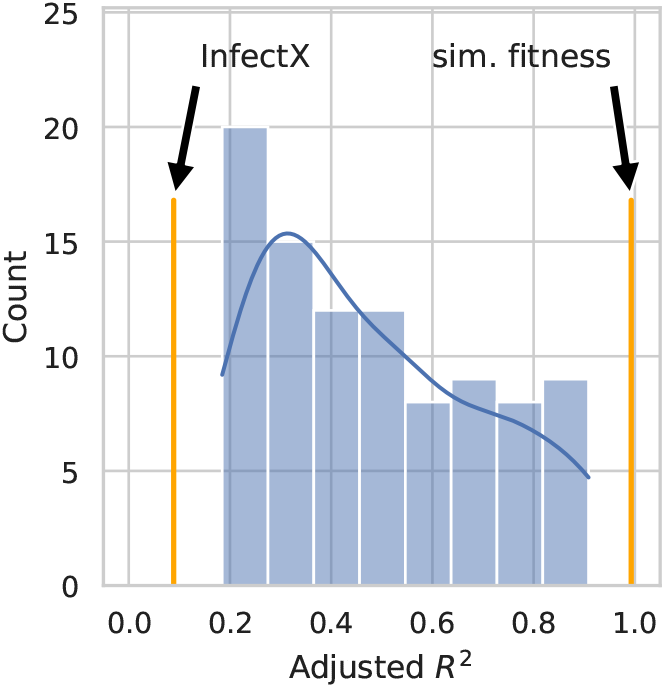
Adjusted *R*^2^ density of simulations in Section 3.1, with additional lines indicating the values using InfectX data, or InfectX with simulated fitness, instead.

### 3.2. InfectX siRNA Data

We run our lasso model on the InfectX data (Section 2.6.1) allowing all pairwise interactions, and halting at λ = 0.05 or the first 2,000 non-zero effects, whichever comes first. Only the genes and gene-pairs with non-zero predicted effects are then included in the matrix **Z**. Last, we fit the phenotype **Y** to this matrix using least-squares regression **Y** ~ **Z**, using these unbiased estimates and p-values as our final result.

We find 26 proposed effects (21 main and 5 interactions) in under two hours. Our method proposes interactions between five genes and TTN, with varying estimated strengths (see Table 2). Two of these interactions, OBSCN and PLEC, are known protein interactions [36].

We find the same set of interactions in repeated runs (bearing in mind that the matrix is shuffled differently each time). This suggests that these are not random choices, but effects strongly supported by the data. The Adjusted *R*^2^ value is only ≈ 0.088, however, indicating that while a better than random fit has been found, the chosen effects do not explain the overall observed fitness particularly well. To investigate this, we consider the difference between fitting two different phenotypes. Firstly, the predicted effects with the measured cell counts from InfectX, and secondly a simulated set that reflects our assumptions.

For the simulation we used the same **X** matrix, but simulated the fitness effects **Y** as a linear combination of randomly chosen gene effects and gene-pair interactions. Every gene had a 10% chance of being assigned an effect, which were sampled from 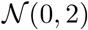. We gave every pair of genes a 0.1% chance of an effect, which were also sampled from 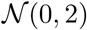. For every row *i* of **X**, the fitness value *y_i_* is the sum of both main and interaction effects present, with additional random noise.

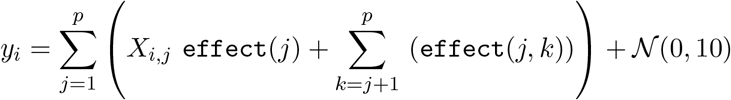

With this simulated phenotype vector, re-running the interaction search with the same parameters, we have an *R*^2^ of ≈ 0.99.

After adjusting for the number of effects proposed, we find that while the fit is better than random using the Z-scored InfectX cell-count as phenotypes, it is not nearly as good as in our simulations. This suggests that at least some of our assumptions are incorrect, namely that our fitness proxy (log cell count) is additive and can be largely explained with individual and pairwise silencing effects, and that the off-target predictions are accurate. While all of these assumptions are somewhat suspect, it should be noted that our siRNA off-target predictions likely miss a significant number of strong effects, and include genes that are not completely silenced [1]. With this in mind, it is plausible that even if the cell count responds to gene silencing according to our assumptions, the predicted effects may not be significantly better than random until accurate siRNA off-target predictions are available.

### 3.3. Antibacterial Resistance

We fit our antibacterial resistance data (see Section 2.6.2) with three different sets of parameters. First, we allow all interactions. Second, we restrict to interactions within 100 columns of each other. Finally, we restrict to interactions within 10 columns of each other. In all cases, we run until the adaptive calibration stopping condition is met. In the first case, allowing all interactions, we find that repeated runs do not suggest any of the same effects. Since the data only contains 259 samples for over 75,000 effects (and over 2.5 billion interactions) it is unsurprising that there are several equally good solutions. We fail to find a reproducible result here because the data simply does not suggest one, and this run is included only to demonstrate that our implementation works at this scale. The second and third cases produce more consistent results, with some common interactions suggested in both cases. While limiting to interactions within 100 columns rules out the majority of possible interactions, it also limits the number possible solutions enough that we can find one reliably with only 259 samples.

Moreover, when we reduce the interaction matrix to only the non-zero predicted effects, and produce an unbiased fit with least-squares regression, we find that our fit explains the variance in resistance extremely well. Restricting interactions to effects within 100 entries of each other, we have a multiple *R*^2^ of 0.99, and an adjusted *R*^2^ of 0.86. Even limiting to interactions within ten entries, we have a multiple *R*^2^ of 0.78, and an adjusted *R*^2^ of 0.63. These suggest that in this case our model is a particularly good fit.

There were 16 sets of variants found in both limited-distance runs (interactions within 10 or 100 columns only). For each of the 16 SNVs their genes, functions, and interactions were assessed. Genes were identified based on PAO1 reference co-ordinates using Artemis [7]. The STRING[36] database was used to assess the validity of the protein-protein interactions.

Five of the SNV pairs occurred in the same gene. There were four pairs that had high interaction scores > 0.7 and two of pairs were identified twice. Many SNV’s were found in genes that encoded for proteins involved in biofilm formation and maintenance indicative of long term chronic infections that are often associated with general antibiotic resistance. Other than pilY1, no other gene was found to be mutated in the lab-based evolution study [33].

There were two pairwise effects that had significant p-values in both runs. The first of these pairs occurred in a gene that encoded a copper resistance protein. The second pair was found in a gene that encodes an RNA binding methyltransferase.

## 4. Discussion

Genotype-phenotype data sets have recently become available at a never before seen scale. In principle, it is possible to infer not only the effect of individual genomic variants within such data, but of pairwise combinations of their effects. While this has been shown to work in theory, and a number of tools have been developed that work on a smaller scale, there is a shortage of effective methods for human genome-scale data. In this paper we present a regression based method for such large-scale inference of pairwise effects.

Our method performs coordinate descent lasso-regression on a matrix containing all pairwise interactions present in the data. For such an approach to work at scale, we had to make a number of improvements. First we parallelised the algorithm by dividing the matrix into shuffled sets for each thread. We then drastically increased the scale of tractable data sets by compressing columns of the matrix using Simple-8b. Combined with the typically sparse binary nature of genotype-phenotype screens, our method is able to effectively consider hundreds of millions of possible interactions.

We compared the accuracy and running time of our work to `glinternet`, the best of the methods we used previously [12], and found that our method provides comparable accuracy and precision while running hundreds of times faster. We also tested our method using two genome-scale real data sets. One is an exome-wide siRNA perturbation screen (*n* ≈ 67,000 siRNAs and *p* ≈ 19,000 genes). The other measures antibacterial resistance with respect to genetic variations in *Pseudomonas aeruginosa*, and includes over two billion possible pairwise interactions. In both cases our method finds a number of effects that are either plausible or previously known.

In some cases we can significantly improve the running time and memory use by only considering local interactions. If interactions are restricted to those within 1, 000 positions of each other, we can search our siRNA screen using ≈ 40GB of memory in ≈ 20 minutes.

While our method is effective on this scale, there are some limitations that would make it difficult to use on significantly larger data sets. Both the time and space requirements are quadratic in the input sequence, and performance does not scale well with non-uniform memory access. This essentially limits our approach to data that fits in memory on a single machine. The pairwise additive model is also something of an oversimplification. It remains unclear to what extent genetic effects be treated as additive, and ignoring interactions among of more than two items could well be leaving out the most important effects. In this case we may end up spuriously associating phenotype changes with individual and pairwise effects that just happen to be present, rather than the true, more complicated, interaction.

There are nonetheless a number of opportunities to expand upon this work. If the original **X** matrix is sparse, and the pairwise interaction matrix **X2** is very sparse, we would expect threeway interaction columns of an **X**_3_ matrix to be even more so. If there are few enough non-zeros in such a matrix, it may be possible to extend our method beyond pairwise interactions without any fundamental changes. While there would be *p*^3^ columns in a three-way interaction matrix, if the vast majority contain only zeros we may still be able to store it. The indices of non-zero three-way interaction columns could themselves be stored in a compressed list of offsets. Any column whose index is not in this list could then be presumed to be zero and left out of beta updates. Since the memory and time requirements only grow with the number of non-zero entries, this could provide a well be enough for sufficiently sparse data.

Alternatively, as we showed in Section 2.5, we can significantly increase the scale of interaction inference methods by reducing the search space. A more targeted approach than restricting the genome distance, estimating distance in 3D space using Hi-C [4] for example, would drastically reduce the time and space requirements, allowing higher order interactions to be considered.

Finally, the interactions proposed in Section 3.2 that have not already been confirmed may well be real, and are worth further investigation.

Our method is implemented in C, and an R package is provided at github.com/bioDS/pint.

## Appendix A Cyclic Linear Regression

As noted in Section 2.1, where *y_i_* is the *i*th element of **Y**, *β_j_* is the *j*th element of *β*, and *X_ij_* is the entry in the matrix **X**_2_ at column *j* of row *i*, the error is the following.

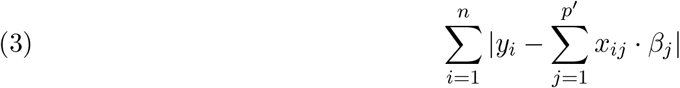

Note that we assume that the fitness vector **Y** is already centred around 0, and omit the offset *u* present in [40]. In the context of pairwise genetic interactions we would rather have a smaller number of definitely relevant effects, than a large number of marginally relevant ones. To this end, we add the lasso penalty to the error in Eq. (1). This penalises large beta values according to a parameter λ, and results in a smaller set of, typically larger, non-zero beta values [37]. With this added penalty we minimise the value:

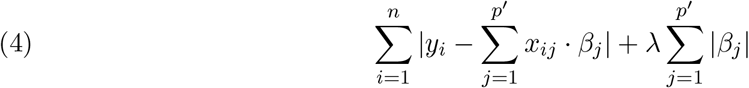

We do this by minimising the component of this error that each *β_j_* is able to account for. For a particular *β_k_*, the component of this error that is affected by changing *β_k_* is:

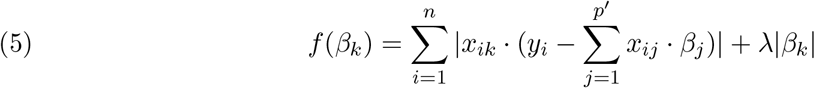

This error comes from the non-zero entries in column *k* of **X**. Since in our case all entries are either 1 or 0, this is simply the sum of errors of rows where column *k* has an entry, with a penalty imposed for large beta values.

To minimise this component *f* (*β_k_*) alone, we define *r_i_* and *S_k_*:

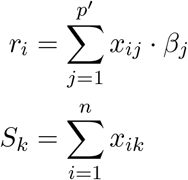

The error affected by a single beta value (Eq. (5)) can then be minimised by updating *β_k_* with Eq. (2), repeated below:

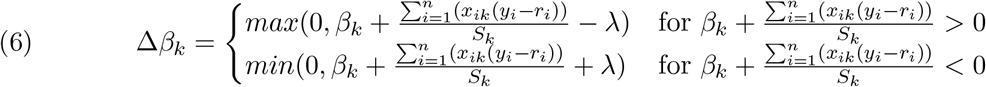

This is equivalent to the solution from [40], as we will now show. Their solution is defined separately for positive and negative *β_k_*:

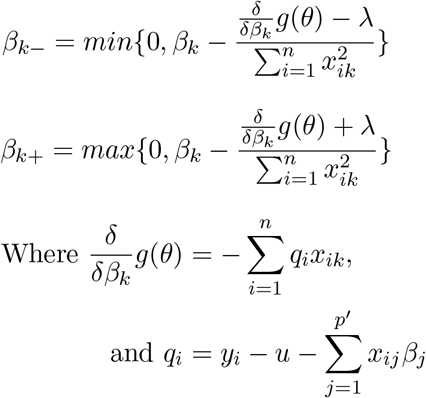

Note that we assume the intercept term *u* = 0, because *Y* is centred around 0, and *u* can therefore be omitted. We shall first focus on proving the equivalence of our construction for *β_k__*. Since *x_ik_* ∈ {0,1}, it follows 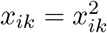, and therefore 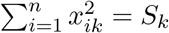. This gives us

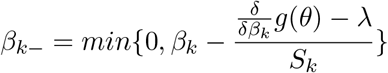

Also substituting 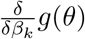, we have:

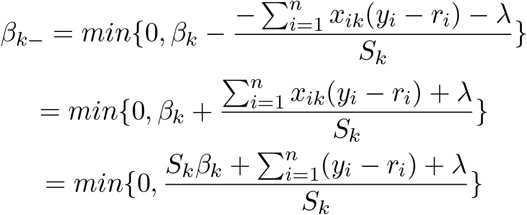

This is equivalent to Eq. (2) for *β_k_* < 0. The positive solution is equivalent, substituting min for max and subtracting λ. Iteratively minimising beta values until the solution converges, we have Algorithm 1. We consider the algorithm to have converged when 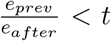 for some threshold *t*, where *e_prev_* is the error before the iteration and *e_after_* the error after the iteration. We arbitrarily chose *t* = 1.0001 as the default in our implementation.

**Algorithm 1:**
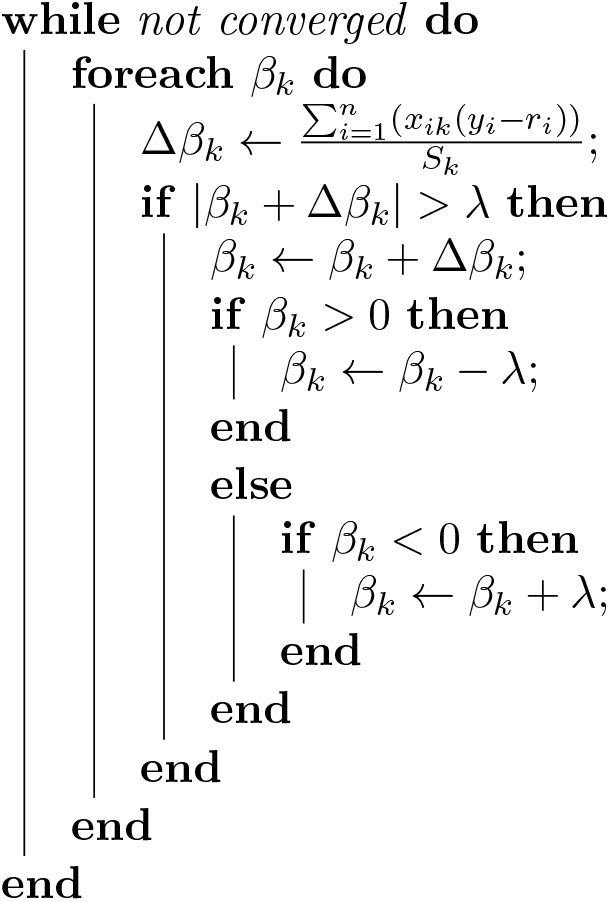
Sequential Cyclic Algorithm

A naive implementation would read every entry of the **X**_2_ matrix, and every value in the vector **Y**, every iteration, for every beta update. With **X**_2_ ∈ {0,1}^*n*×*p*’^, 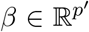 and 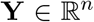, this is Θ(*np*^4^) operations per iteration. Since *x_ij_* and *y_i_* are constant, 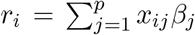 only changes when *β_j_* changes. Updating this value every iteration, rather than re-calculating it, we only need to perform n operations per *β* update. This brings the number of operations for an iteration down to *np*^2^. To update *β_j_*, we now read a single column, *X_j_*, and the values of *r_i_* and *Y_i_* for each non-zero entry *x_ij_*. By including **Y** in **R** such that 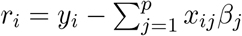 for all *i*, we no longer need to read **Y**.

We can further reduce the work that needs to be done by storing a sparse representation of **X**_2_. While **X** is a sparse matrix, **X**_2_ is an extremely sparse matrix. In a typical simulated data set from [12] we go from, on average, 112 out of 1, 000 entries per column in the **X** matrix to 16 out of 1, 000 in **X**_2_. We therefore store **X**_2_ in the following format. Each column is a list of the positions of its non-zero entries (Fig. 7). Since these are one by definition, we don’t store their value. We store the matrix column-wise to ensure the column *X_j_* can be read quickly when updating *β_j_*. Each column is therefore stored as a separate array of integers.

**Figure 7.**
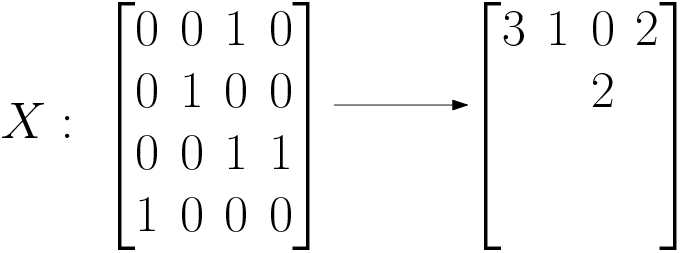
Simple sparse matrix representation

## Appendix B Compression

We can considerably reduce the size of this matrix in Fig. 7 by compression the columns. Since we have a sequence of increasing integers we can store only the offset from the previous entry, keeping the entries small. The resulting sequence of (mostly) small numbers can then be efficiently stored using integer compression methods. We describe the compression method we use in Appendix B.1 and compare it to other methods in Appendix B.2.

### B.1. Simple-8b

Simple-8b is a non-SIMD compression scheme, with performance comparable to other state of the art methods [35, 25, 38]. While SIMD-based compression schemes can often offer significantly improved compression and decompression speed [22] [35], their implementation is architecture dependant. Simple-8b only requires a CPU be able to efficiently handle 64-bit arithmetic, and does not significantly underperform compared to state-of-the-art SIMD techniques in our testing (Appendix B.2).

Simple-8b is a 64-bit variation of the Simple-9 encoding scheme [2], and stores a sequence of integers in a single 64 bit word. The number of integers stored depends on the size of the largest one, and is indicated by a four bit ‘selector’. The remaining 60 bits are divided into integers of size 1, 2, 3, 4, 5, 6, 7, 8,10,12,15, 20, 30 or 60, with between 240 (only possible if all values are zero) and one integer stored. As seen in Fig. 8, this considerably reduces the size of **X**_2_ in our test data (two sets from [12], one with *p* = 100, *n* = 1, 000, another with *p* = 1, 000, *n* = 10, 000). In the larger *p* = 1, 000 set, total memory use is reduced by over 85% compared to storing integers directly. It is worth noting that this compression works well even for non-sparse sections of the matrix, since the offsets are extremely small. In an extreme case, we can store up to 240 sequential 1’s in a single 64-bit word. While the earlier offset-based format (Fig. 7) relies on the sparsity of the matrix for its efficiency, compression works well regardless.

**Figure 8.**
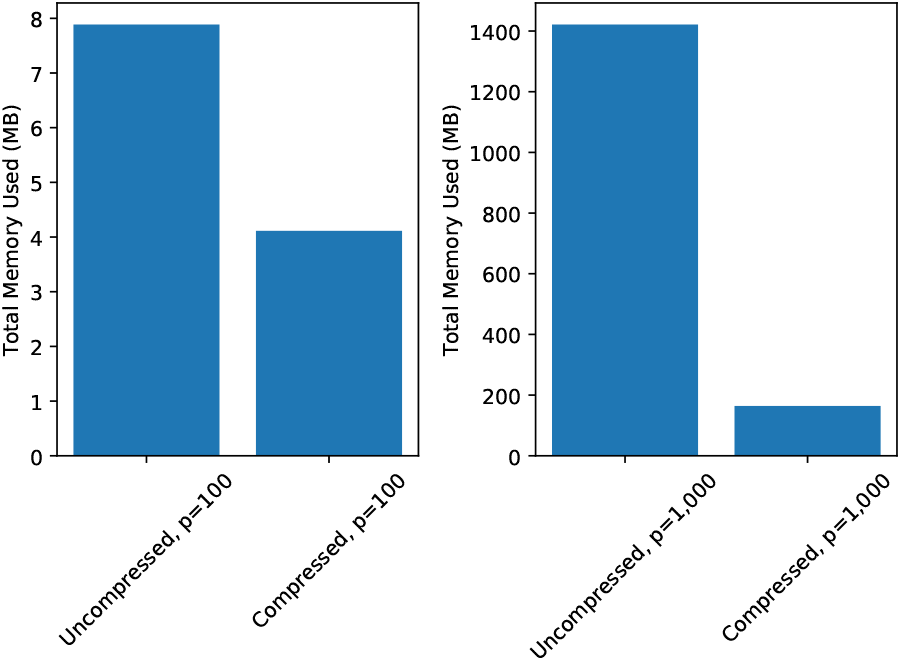
Compression effect on memory use. Note that this is the total peak memory use of the program, not solely the memory used by the matrix **X**_2_. In both cases *n* = 10 ·*p*.

### B.2. Comparing Methods

While Simple-8b allows our implementation to be used on any 64-bit CPU, we could also take advantage of SIMD-based methods where the such CPU instructions are available. To determine whether this is a worthwhile improvement, we compare our Simple-8b implementation to a number of state of the art alternatives.

Recent work suggests TurboPFor [29] has a particularly high compression ratio [38]. We therefore compare the best performing methods from TurboPFor against our implementation of Simple-8b (Fig. 9). The tests are performed using 32 threads across two eight-core (16 SMT threads) Intel(R) Xeon(R) Gold 6244 CPUs in a NUMA system. To compare these methods, we perform 50 regression iterations on a test data set of *p* = 1, 000 genes and *n* = 10, 000 siRNAs. We examine the total time taken for the process, as well as the total memory used and time for the regression function alone (excluding calculating and compressing the interaction matrix).

**Figure 9.**
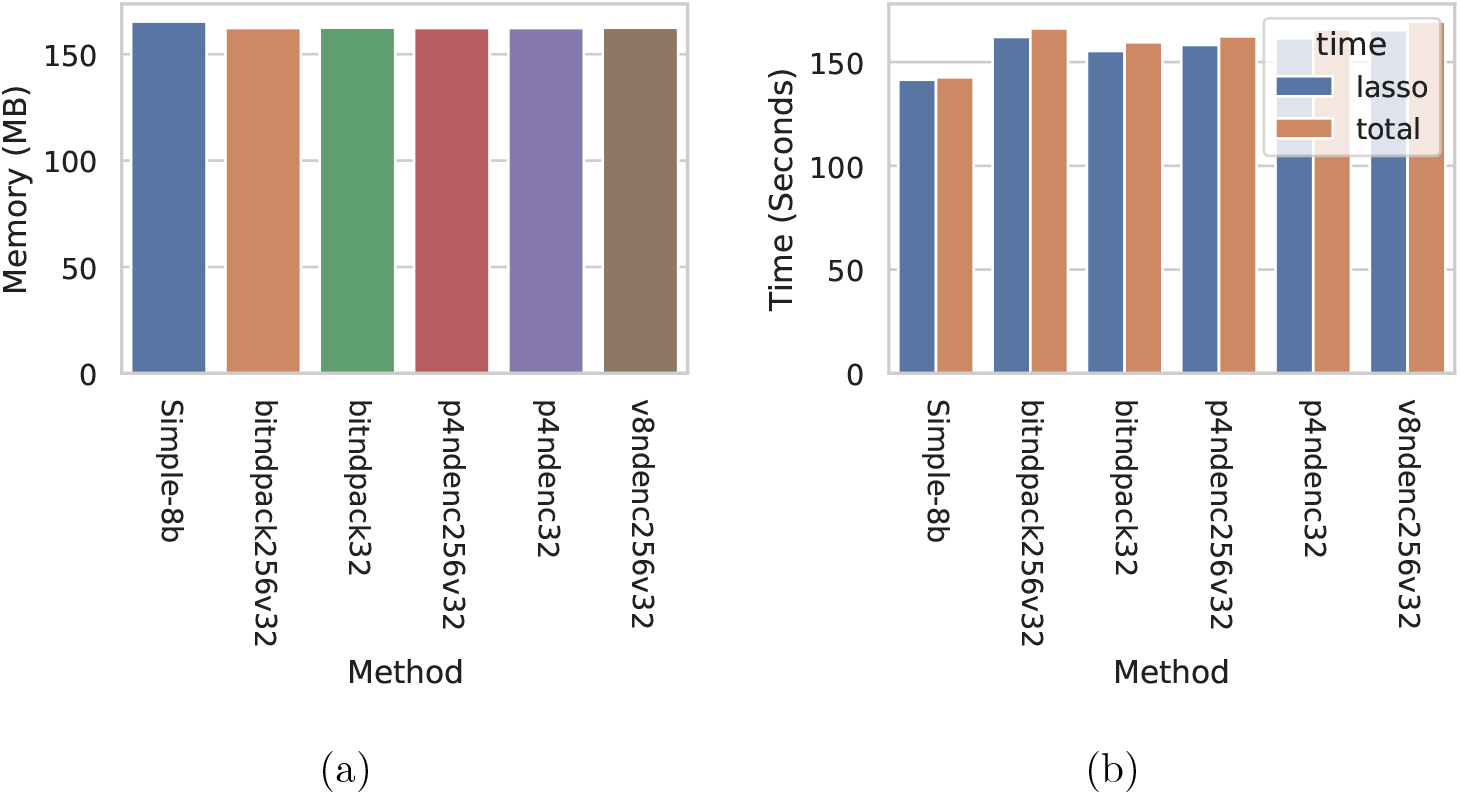
Comparison of Compression Methods. (a) Total memory used, compressing the sparse **X**_2_ matrix with each method. (b) Total time taken and time taken (including compressing **X**_2_) and time taken for lasso regression alone, using each method.

We see that both the time to produce the compressed matrix (seen in Fig. 9 as the difference between total time and lasso-only time), and the running time are comparable for all TurboPFor methods.^2^ While every TurboPFor method we tested improved the compression ratio compared to Simple-8b (Fig. 9a), we consistently found that the running time was longer (Fig. 9b). It is possible that this is a result of the way the columns are being read in each method. Using TurboPFor, we compress and decompress entire columns at a time. With our Simple-8b implementation, we process each 64-bit word separately. This allows us to use the column as it is being decompressed. Avoiding re-reading the column after decompression also allows the entries to be evicted from the cache earlier.

While it is also possible to process compressed words as they are read using the tested TurboPFor methods, there does not appear to be a significant difference in compression.

## Appendix C Choosing Lambda

The regression parameter lambda determines how large a change to β needs to be before it will actually be made. During Lasso-regression we begin with an extremely large value of lambda, and gradually decrease this until we think smaller effects are only going to over-fit the data. For a better estimate of effect strengths, we only use lasso regression to choose the non-zero beta values, and then perform Ordinary Least Squares (OLS) regression on the non-zero entries. The *β_i_* estimates from OLS regression are the effect strength estimates and the p-values are used to determine whether an effect is significant. It suffices then to continue decreasing lambda until an arbitrary small lambda value, relying on OLS regression to filter for significant results. As lambda decreases the number of non-zero effects increases, however, eventually becoming too many for OLS regression. Since small effects are less likely to be correct [12], we prefer to stop at a larger lambda.

We provide two options for choosing the final lambda in our package. First, we choose lambda such that the number of non-zero effects is small enough for OLS regression. Second, we implement a fast method for empirically choosing a reasonable stopping point.

In either case we begin with lambda sufficiently large that all beta values will be zero. Lambda is then gradually decreased, setting the new value at each step to *λ_new_* = 0.9o *λ_prev_*. We decrease lambda until we reach or pass the minimum value (0.05 by default). After fitting with each lambda, we optionally check one (or both) of the two stopping conditions. First, we can check whether we have reached the maximum number of non-zero beta values. Alternatively, we perform the adaptive calibration test [8], stopping if the conditions are met.

We use the adaptive calibration lambda selection method from [8] instead of the standard *K*-fold cross-validation because cross-validation requires fitting each lambda value *K* times, and this increase to the run time is unacceptable for large data. Adaptive calibration only requires a single relatively small calculation for each lambda. It aims to choose the minimum value of lambda that is sufficient to control fluctuations. Assuming **X**_2_ satisfies the design condition from [39], the value chosen is within a constant factor of this ideal value, and precision and recall are comparable to cross-validation [8].

In Fig. 10 we compare precision, recall, and running time when using the adaptive calibration stopping condition to running as many iterations as we can, on a simulated data set from [12]. In one case we decrease lambda until the adaptive calibration condition is met. In the other we limit the number of non-zero effects to 2, 000. In both cases, we then perform the OLS regression step and filter out results with a p-value ≥ 0.05.

**Figure 10.**
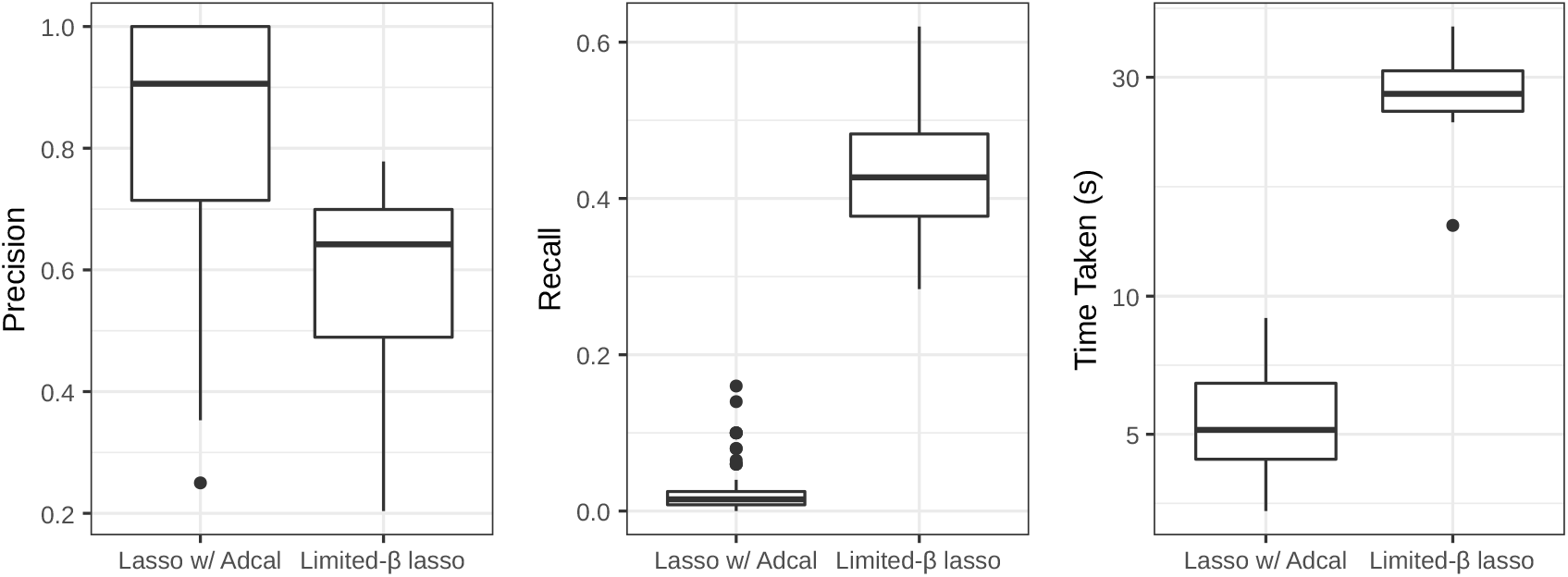
Adaptive calibration effect on large sets. ‘Limited-β Lasso’ runs until 2, 000 beta values are non-zero, continuing until the cutoff otherwise. ‘Lasso w/ Adcal’ halts once adaptive calibration conditions are met, continuing until the cutoff otherwise.

We see significantly higher recall when running until a very small cutoff, allowing only 2, 000 effects in total to allow OLS regression. Adaptive calibration on the other hand stops very early. While the predictions at this point are almost entirely correct, the majority of the effects that would be found with lower lambda values are missed. Since the loss of precision at lower lambda values is mitigated effectively by OLS p-value filtering, we suggest this approach in general.

Both the adaptive calibration option and limiting non-zero betas can prevent the algorithm from finding small effects. This is in many cases beneficial, as it is the small values of lambda, and therefore the small effects, that are computationally expensive. Moreover, our previous benchmarks [12] show that small effects are also the least likely to be correctly identified. With that in mind, we do not consider ignoring these effects to be a problem.

## Appendix D Parallelisation Details

In this section we provide an overview of the challenges in parallelising Lasso-regression, and our attempts to overcome them. Appendices D.1 to D.3 describe in detail the way effect strengths (beta values) are updated, and the barriers to doing so in parallel. Appendix D.4 covers the methods we investigated, but do not use in our final implementation. The shuffled method we finally used is described in detail in Appendix D.5.

### D.1. Beta updates

We refer to an update of a βj corresponding to column *X_j_* as a column update. Within a column update, there are no barriers to running in parallel (i.e. parallelising over rows). We can iterate through the elements of the column in parallel using openMP, and calculate the sums with a reduction. The contents of a single column are stored sequentially in memory, which limits the effectiveness of such an approach. The contents of the columns are only read, and not written, in this process, so there is no overhead in maintaining cache coherency. Once a single value has been read on one core, an entire cache-line will be available from its local cache, however. Since these have been read from memory already, there is no advantage to reading them into another core’s cache for parallel processing. We could attempt to offset the work of each core, so that each will be working on a separate cache line within the same column of the matrix. Such an approach, however, assumes that the column contains at least 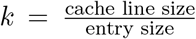. (num cores) entries, which is unlikely. There is also considerable overhead in thread barriers [26], and the work done must be enough to justify this. To solve these problems, and avoid having threads idle when their component of the work is finished, we would need to have several times k entries. In our test set of *n* = 10,000 siRNAs, the mean number of non-zero entries in a column is only 150, or seven compressed 64-bit words. The L1 data cache of our test CPU (A Xeon Gold 6244) has a 64 byte cache line size, enough for eight entries. Even with hundreds of thousands of siRNAs, each column could only be expected to be a few cache lines. Running the iterations over columns in parallel, rather than rows, is therefore the focus of our parallelisation attempts.

### D.2. Overlap Error

We cannot simply perform several column updates in parallel. Each column update both reads and writes *r_i_* values for every non-zero entry in the column. If two columns are updated in parallel, and they both have non-zero entries in a common row, there is a time-of-check to time-of-use problem. An update can occur in an *r_i_* after that value has been read by another thread. While the cached *r_i_* themselves will never be incorrect, as the updates are atomic and always the result of real changes to a beta value, both columns will be updated based on the old value. Both columns are partially responsible for the difference between the current fit, 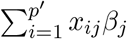, and observed fitness, *y_i_*, of this common row. Each of these updates will attempt to minimise this as much as possible, without taking the other update into account. This can result in overcorrecting for the error in the common row, potentially increasing the overall error. To compensate for the increased error, the next update may make an even larger change to *β* (Fig. 11a). In the worst case, if two or more updates repeatedly overcorrect for each other, this can prevent convergence entirely (Fig. 11b).

**Figure 11.**
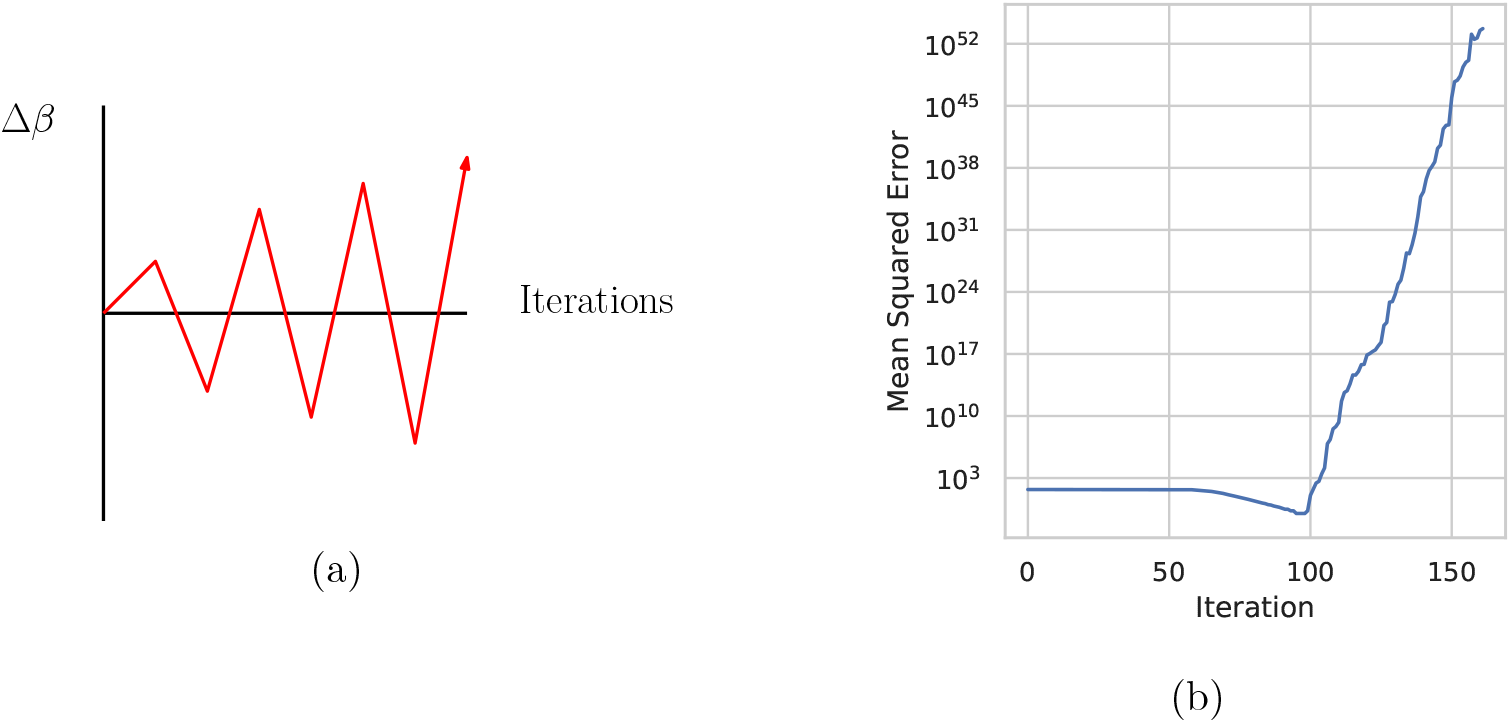
Effect of repeated overcorrection. (a) Repeated overcorrections lead to increasingly large changes to error. (b) Effect of repeated overcorrection on fit. This example is the result of fitting a 50 × 20 random binary **X**_2_ matrix, with random **Y** values between −1 and 1.

### D.3. Deriving Overcorrection Error

Simultaneous updates may result in overcorrection, but we can analytically determine exactly how much. Using the definitions from Appendix A, ignoring the lasso penalty for the moment, we update a single *β_k_* by Δ*β_k_* as follows.

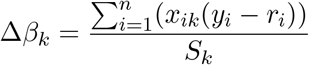

Let us define 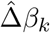 to be the value that Δ*β_k_* would take if, for every *j* < *k*, the update to *β_j_*, had already been performed. If we perform all updates strictly sequentially, then 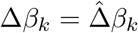. Similarly, we define 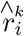 to be the value of *r_i_* after all updates prior to *k* have been performed.

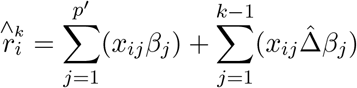

Since the only difference between 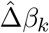 and Δ*β_k_* is the change in *r_i_* caused by previous beta updates, it follows:

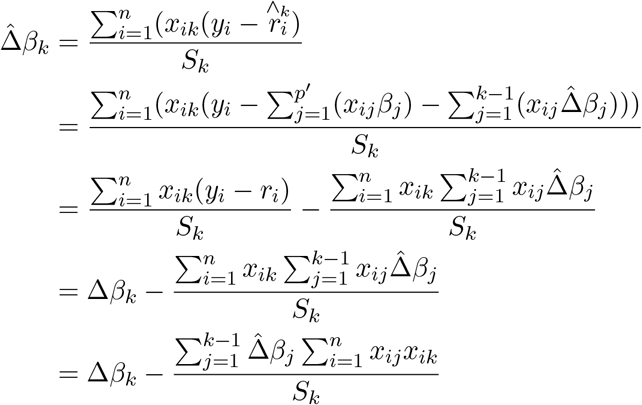

Note that 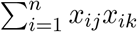 is constant with respect to changes in *β* and **R**. We can compute these values once after the input has been read, and re-use them in every iteration. If we define the overlap between columns j and k of **X**_2_ to be 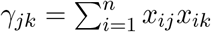, we have the following.

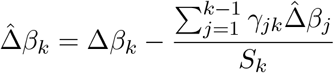

*Remark* 1. 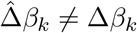 if and only if *γ_jk_* ≠ 0 for some j < k.

For λ ≠ 0 we define the soft threshold function f_λ_(*x*).

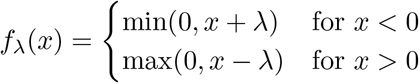

We find that the value of 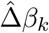 for 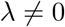 is the following, and use this definition in our package.

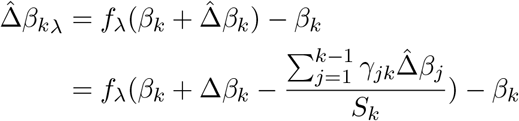

We now consider a concrete example of Remark 1. Let each residual *r_i_* be fixed (for the moment), and update first *β*_1_, then *β*_2_, the intended sequential update effect is then:

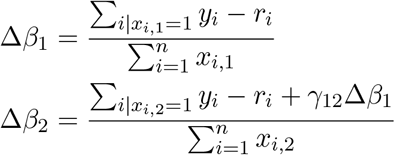

When both updates are instead performed at the same time we get:

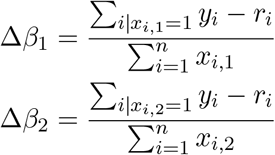

Updating in parallel, the effect of *β*_2_ is estimated based on the original **R**, rather than those that account for the changes made to β**1**. We can easily calculate the expected overcorrection in this case, *γ*_1;2_Δ*β*_1_. Note that we are atomically updating the residuals *r_i_* that are affected by both updates. If we fail to do this, overcorrection becomes difficult to predict.

For example, *β_1_* and *β_2_* correspond to the columns in Fig. 12a, and suppose these columns of **X**_2_ are chosen for simultaneous updates. We can calculate the changes Δ*β_1_* and Δ*β_2_* in parallel and update the residuals safely, because there are no shared values being updated by both threads. In Fig. 12b, we find that both the update to *β_1_* and the update to *β_2_* affect residual *r*_3_. While atomic updates to the actual value of *r*_3_ will guarantee that we finish with the value *r*_3_ + Δ*β*_1_ + Δ*β*_2_, we have not taken the changed value of *r*_3_ into account when calculating Δ*β_2_*. The correct update would have been 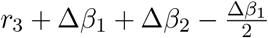

**Figure 12.**
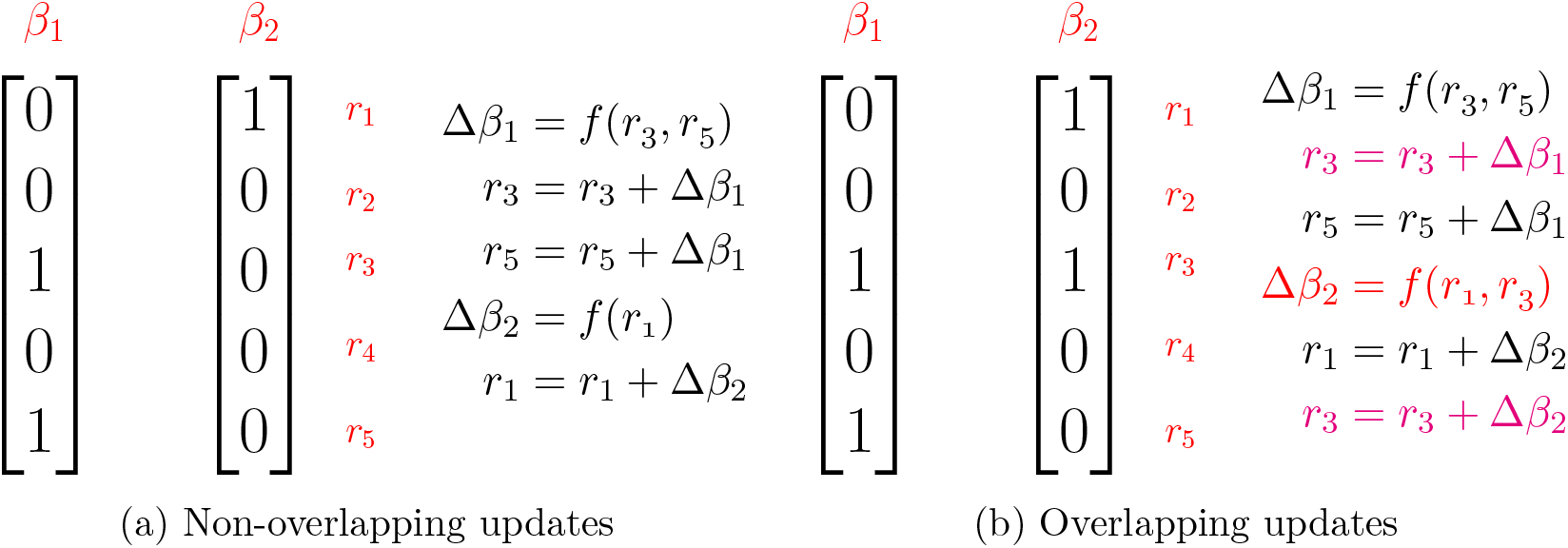

Given that we can safely perform updates for columns that have no overlap, and we can explicitly compensate for the error of sets of columns, we investigate three approaches for parallelisation: compensating for the error of pre-determined sets (Appendix D.4.1), simultaneously updating non-overlapping sets (Appendix D.4.5), and randomly updating shuffled columns (Appendix D.5).

### D.4. Alternative Parallelisation Methods

Two alternative approaches to parallelisation were considered, but not used in the final implementation. In this section we provide a detailed explanation of the barriers to parallelised lasso regression and the methods we attempted, but did not use.

#### D.4.1. Explicit error compensation

To update *β_a_* and *β_b_* at the same time, we need to subtract the overcorrection from one of them (arbitrarily chosen) afterwards. The final value is then the same as if we had updated the values sequentially. Subtracting the overcorrection from Δ*β_a_* we update βa as follows:

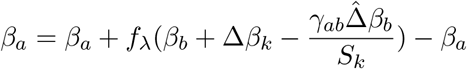

Similarly, we can update any subset of the beta values {*β*_1_,…,*β_p′_*} simultaneously, as long as we account for overcorrection in each update. For every *β_k_* in the subset {*β*_1_,…, *β_l_*}, we make the following correction:

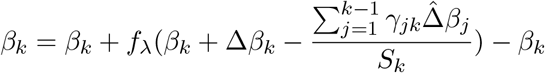

This method has been implemented in the error_comp branch of our repository^3^, and is marginally faster than sequential updating with the right parameters. It does not scale well enough that we can recommend its use, however.

To understand the scalability of this approach, we begin by noting that the time taken to correct *C* simultaneous beta updates is on the order of *C*^2^. This is because each update 0 ≤ *i* ≤ *C* requires reading the *i* − 1 previous corrected values, resulting in 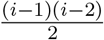 reads. If we were to attempt to update the entire interaction matrix in parallel, followed by correcting errors, there would be on the order of *p*′^2^ corrections. Since a sequential iteration requires only *p′μ* steps, where *μ* is the mean number of non-zero entries per column, we would spend more time on corrections than updates. Even if corrections were run in parallel, this would be slower than sequential updates.

We do not, however, have to update the entire matrix at once. If we restrict ourselves to updating small sets, where ‘small’ is some function of the number of threads we are able to effectively use, the problem becomes tractable. Performing C parallel updates, where C is some constant multiple of the number of available threads, we have in total 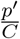 sets to update, resulting in 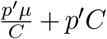 update operations on the main thread.^4^ For 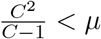 this is an improvement, even with single-threaded correction.^5^

*Remark 2.* Ahmdahl’s Law implies a best-case improvement of:

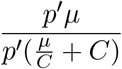

#### D.4.2. Parallel correction does not scale

To improve upon Remark 2, we have to correct beta updates in parallel. This requires calculating 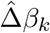 without using any previously corrected values, 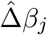 for *j* < *k*. Substituting 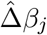, a corrected update 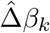 then becomes the following:

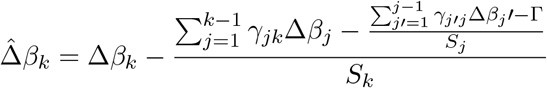

Where Γ is a nested sum containing a further (*k* − 4)! additions. Even for very small sets, computing this is not feasible. We must therefore fix overcorrection sequentially, and accept the limit in Remark 2.

#### D.4.3. OpenMP barrier overhead requires large C

To achieve the improvement in Remark 2, we would need a set of eight columns, updated on eight CPUs, to finish eight times faster. This is in practice not the case. We demonstrate this by running a single iteration on the same test set with varying parallel update set sizes *C*. Using sets of size m times the number of available cores we gradually increase m and measure the time taken to perform all *p′* column updates, without compensating for overcorrection. As we can see in Fig. 13, we require blocks of 64 to 128 times as many columns as there are cores to achieve speed-ups close to the theoretical limit, and at least eight times as many for any significant improvement.

**Figure 13.**
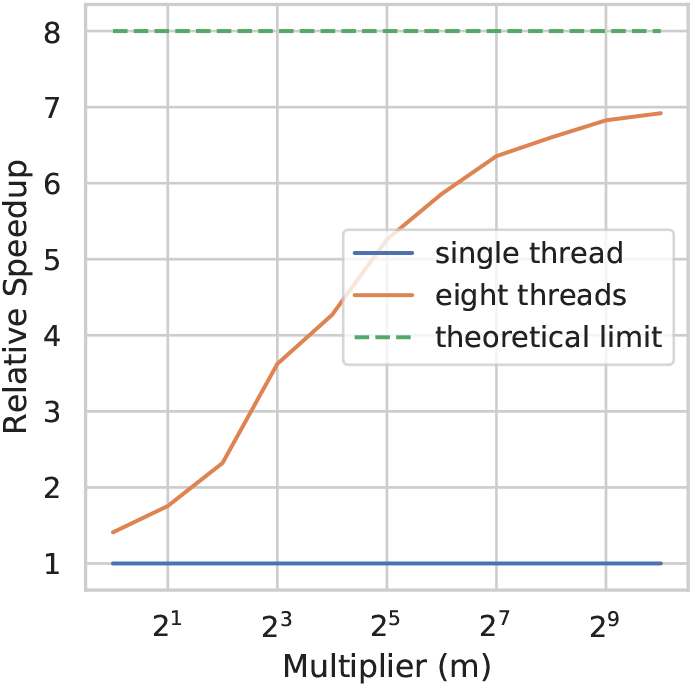
Set size multiplier effect on parallelisation speed-up. Sets contain (number of cores) × C columns.

We find that for small block sizes, the majority of the time all threads are handling an openMP barrier at the end of a parallelised loop.^6^ There are two likely causes. First, when running a small number of column updates in parallel, some of these will often finish before others. If there are no further columns in the set, the thread will then have to wait for the set to finish updating. Secondly, updating columns with entries in nearby rows will mean accessing nearby memory locations to update the residuals. Since these are both being read and written to, maintaining cache coherency across CPU cores is likely to affect performance. The barrier at the end of each parallel section may also take a significant amount of CPU time, as it requires some communication between threads.

Memory access is also less efficient with multiple threads. When a single thread performs a sequence of column updates, these columns are stored in memory sequentially, and each column may be as little as a single word. The core will read in an entire cache-line at once, and this may contain part of the following column, if not the following several. In this case, additional updates for these columns may be performed without any extra memory reads. If, on the other hand, we have eight columns shared between eight threads, they will each still need to read in these values. In our case the eight cores have a shared L3 cache, individual L1 and L2 caches, and 64 byte cache lines. Since one Simple-8b word is 64 bits, we have eight such words in a single cache line. In the worst case, this is eight separate columns. Supposing this is the case, a single read from memory will bring all eight columns into the shared L3 cache. It will then take eight separate reads into various L1 caches, for eight separate cores to perform updates for these columns, whereas a single core would require only one read from L3. With larger data sets, and hence larger columns, this becomes less of a problem. The fact remains though, that cache use is significantly better when each core updates several sequential columns.

All of these issues are mitigated by increasing the size of sets, and updating several sequential columns on each thread. Fig. 13 suggests sets should be at least sixteen times the number of available threads in size. This increases the time required to compensate for overcorrection.

**Figure 14.**
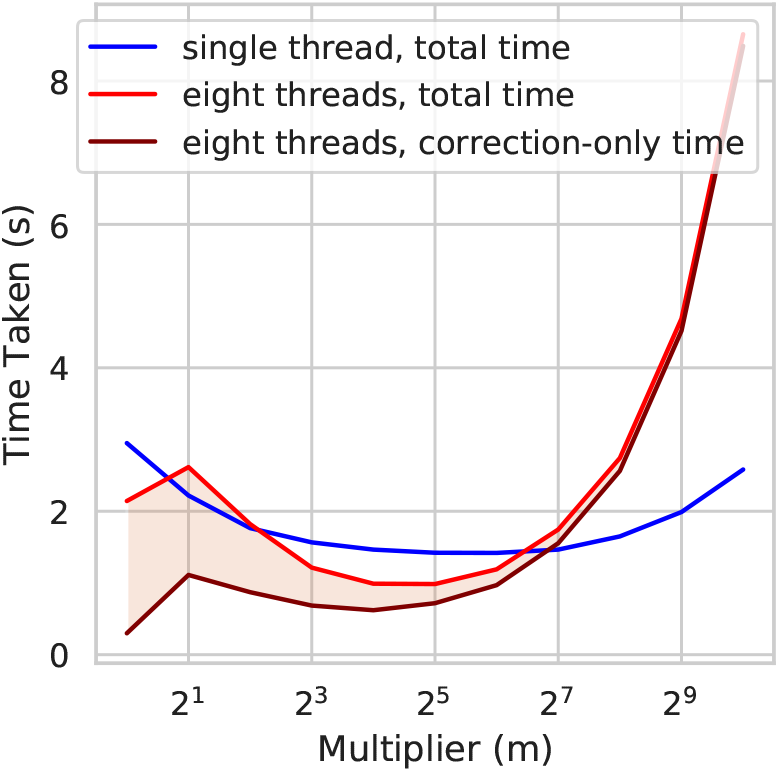
Total time taken (in seconds) for various block size multipliers. Time spent in parallel section is highlighted. Single-threaded time is included for comparison.

#### D.4.4. There is no effective value of C

The results in Appendix D.4.2 require us to perform error correction sequentially, i.e. on a single core. As we saw in Remark 2, for sufficiently small sets this is not a serious limitation. However Appendix D.4.3 also suggests that we may need large sets.

We first note that we can’t significantly reduce the time taken in the error correction step. Updating **R** and calculating corrections take approximately half the time each. While updating **R** can be parallelised, there is so little work here that the thread barriers again result in worse performance. In the overlap matrix, containing the overlap between columns in the set (*γ_ij_* for columns *i, j*), around 50% of the entries are non-zero. Removing these or using a sparse compressed matrix to store overlap is therefore unlikely to be a significant improvement.

Fig. 14 shows the amount of time spent in error correction increases quadratically with the block size. At a multiplier of 256 this overtakes the entire update time on a single core, and we see that the multi-threaded implementation becomes slower than the single-threaded. The best multi-threaded result we see is at a multiplier of thirty-two, where the parallel version achieves a 44% improvement in iterations per second. Even with improvements to the error correction routine, it is unlikely that this approach to parallelisation can achieve more than double the performance of the sequential version. We therefore arrive at the conclusion:

*Remark* 3. Parallel updates of sets, followed by error correction on those sets, is not a feasible approach for parallelisation.

#### D.4.5. Limiting overlap

While we cannot compensate for overcorrection after the fact, Fig. 13 nonetheless suggests that there is some hope for performing a block of updates in parallel. Rather than allowing these blocks to be arbitrary, we now consider restricting updates to blocks of non-overlapping columns of **X**_2_. Again, we first divide the matrix into sets of columns for which we can perform updates simultaneously, then update these sets one at a time. Here, these sets will be collections of columns that either do not overlap, or overlap very little.

#### D.4.6. Sets of no overlap

Since the columns of **X** are relatively sparse, and the columns of **X**_2_ particularly so, it is plausible that we could find sets of a few non-overlapping columns purely by chance. In our test set of 1, 000 columns and 10, 000 rows the mean number of non-zero entries in a column is 0.58%. In this case, we would expect the fraction of entries in common between two randomly chosen columns to be (0.58%)^2^, or 0.34 entries per column. If we choose two random columns we can reasonably expect them not to overlap. Extending this we find sets of non-overlapping columns using the following method.

We start by randomly shuffling the columns. We then compare every second column with its neighbour, recording a set of two non-overlapping columns where possible (Fig. 15b). Smaller sets are then repeatedly merged to form larger sets, where no overlap exists between columns (Algorithm 2). The algorithm described in Algorithm 1 is then run, updating beta in parallel for each set of non overlapping columns.

**Figure 15.**
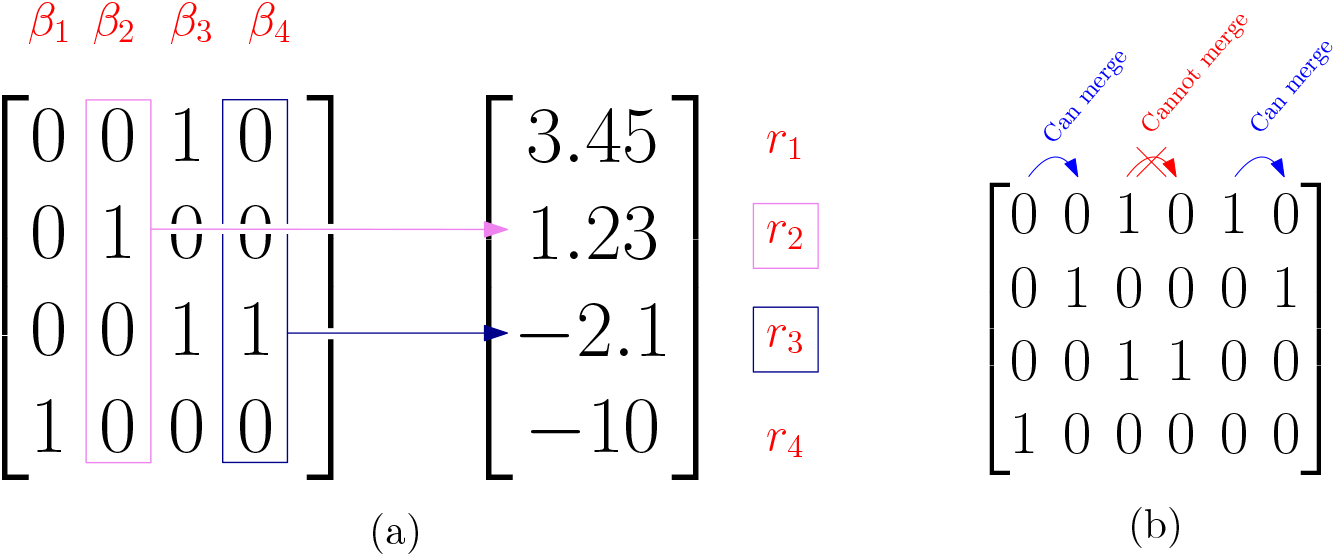
(a) Completely non-overlapping updates affect different residual values and can be done safely in parallel. (b) Attempting to merge neighbouring columns.

**Algorithm 2:**
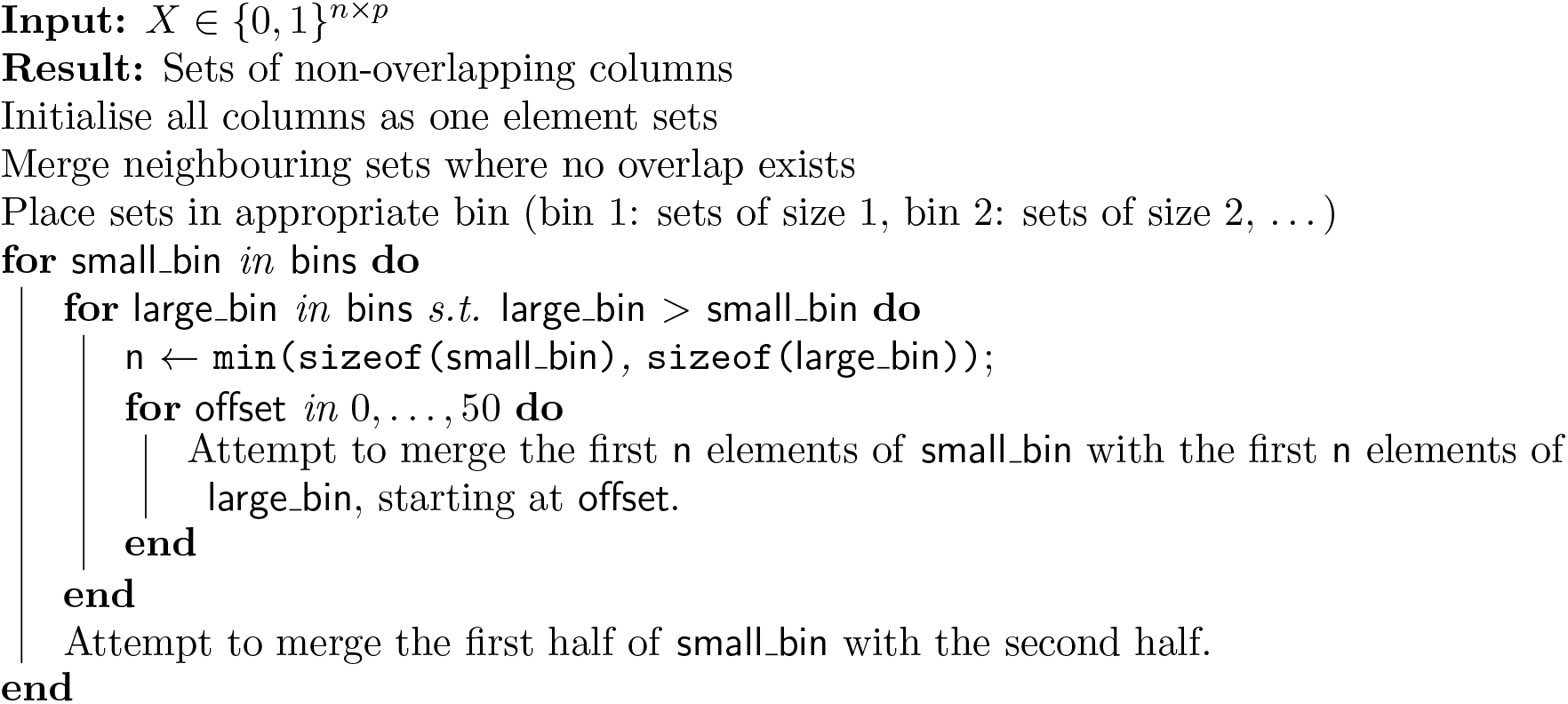
Mergesets Algorithm

#### D.4.7. Performance of Overlap Method

We compare run time on one and four cores, using simulated sets of 1000 siRNAs and 100 genes (i.e. pairwise interactions on a 1000 × 100 matrix) in Fig. 16. Once sets of non-overlapping columns have been found, updating non-overlapping sets in parallel improves run time compared to the single-threaded version (Fig. 16b).

**Figure 16.**
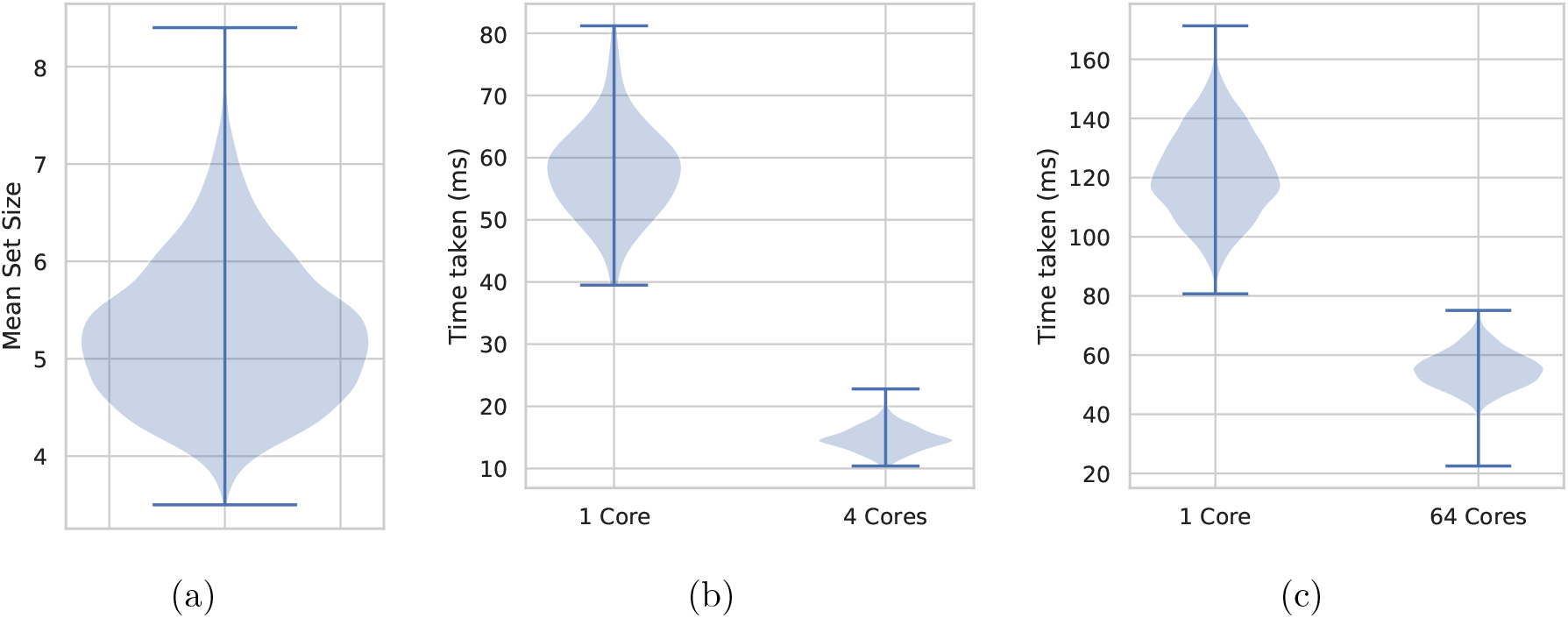
(a) Mean non-overlapping set sizes. (b) Run time after finding mergesets on four vs one core. (c) Run time after finding mergesets on 64 vs one core. Note that the 64 core system has considerably slower memory than the four core system.

As the size of input data increases, however, finding sets with absolutely no overlap becomes more difficult. Since, as explained in Appendix D.4.1, efficiently using more threads requires larger set sizes than those in Fig. 16a, we aim to improve on these. Running more comparisons with our current implementation is not feasible, finding sets already takes as long as the regression step. Instead, we relax the criteria from no overlap to very little overlap.

#### D.4.8. Partial Overlap

We can find much larger sets of simultaneously updateable columns if we allow a small amount of overlap between these columns. While this does allow some error to be introduced in the calculation of beta updates, we expect that by limiting the overlap this error will remain small, and not result in the drastic overcorrection seen in Fig. 11.

In small test sets (n = 1000, *p* = 100), increasing the available overlap significantly increases the found set size (Fig. 17a), also improving the run time. Interestingly, increasing the allowed overlap to 100% does not harm the run time (Fig. 17b) or the mean squared error of the final fit (Fig. 17c). We would expect both of these to suffer when the columns significantly overlap, as either further iterations are required to correct for the introduced error, or overcorrection error is allowed to remain. It appears that even allowing 100% overlap, there is a negligible amount of overcorrection occurring.

**Figure 17.**
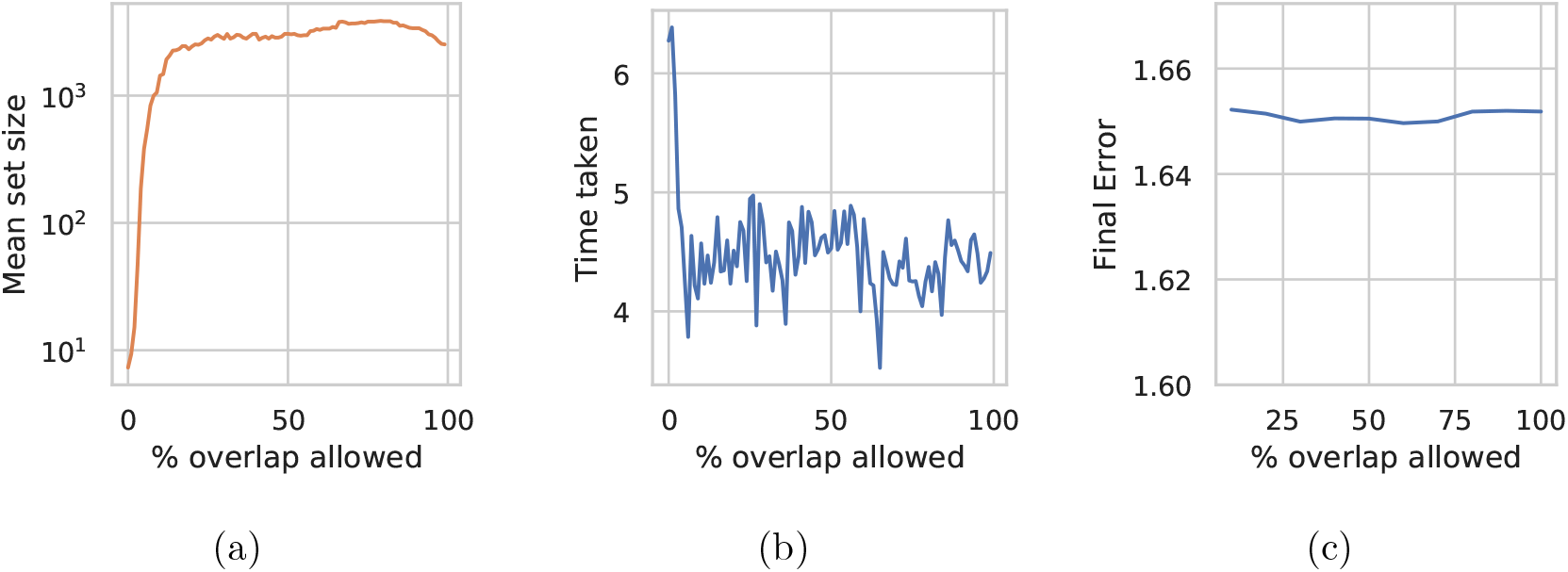
(a) Mean set size, (b) run time, and (c) final mean squared error on 64 cores as allowed overlap increases from 1% to 100%.

### D.5. Random Overlap

Despite the possibility of over-correction, simply updating columns in parallel often works in practice. In fact, as long as we do not update the same columns at the same time every iteration, over-correction does not occur. To avoid this we shuffle the columns every iteration (see Appendix D.4 for a discussion of alternative methods). This was shown to work by Bradley et al. [6] for their lasso implementation, with up to 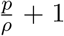 parallel updates, where ρ is the spectral radius of **X**^T^**X**, so long as the columns being updated simultaneously were chosen at random. In our case, using the matrix of pairwise interactions, this allows 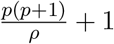 parallel updates, where *ρ* is the magnitude of the largest eigenvalue of 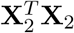. In our smallest test case (*n* = 1,000, *p* = 100), this would allow 222 simultaneous updates. This number increases with larger input data. Combining this with the lambda sequence from Section 2.2, we have our final lasso algorithm, Algorithm 3 (see Appendix A for the full details of the nonparallel algorithm). In this example we have two threads, each working on three columns. Both threads shuffle their columns, then iteratively calculate changes and update each columns β value. Note that this calculation requires reading all β values, including those being updated by the other thread.

**Algorithm 3:**
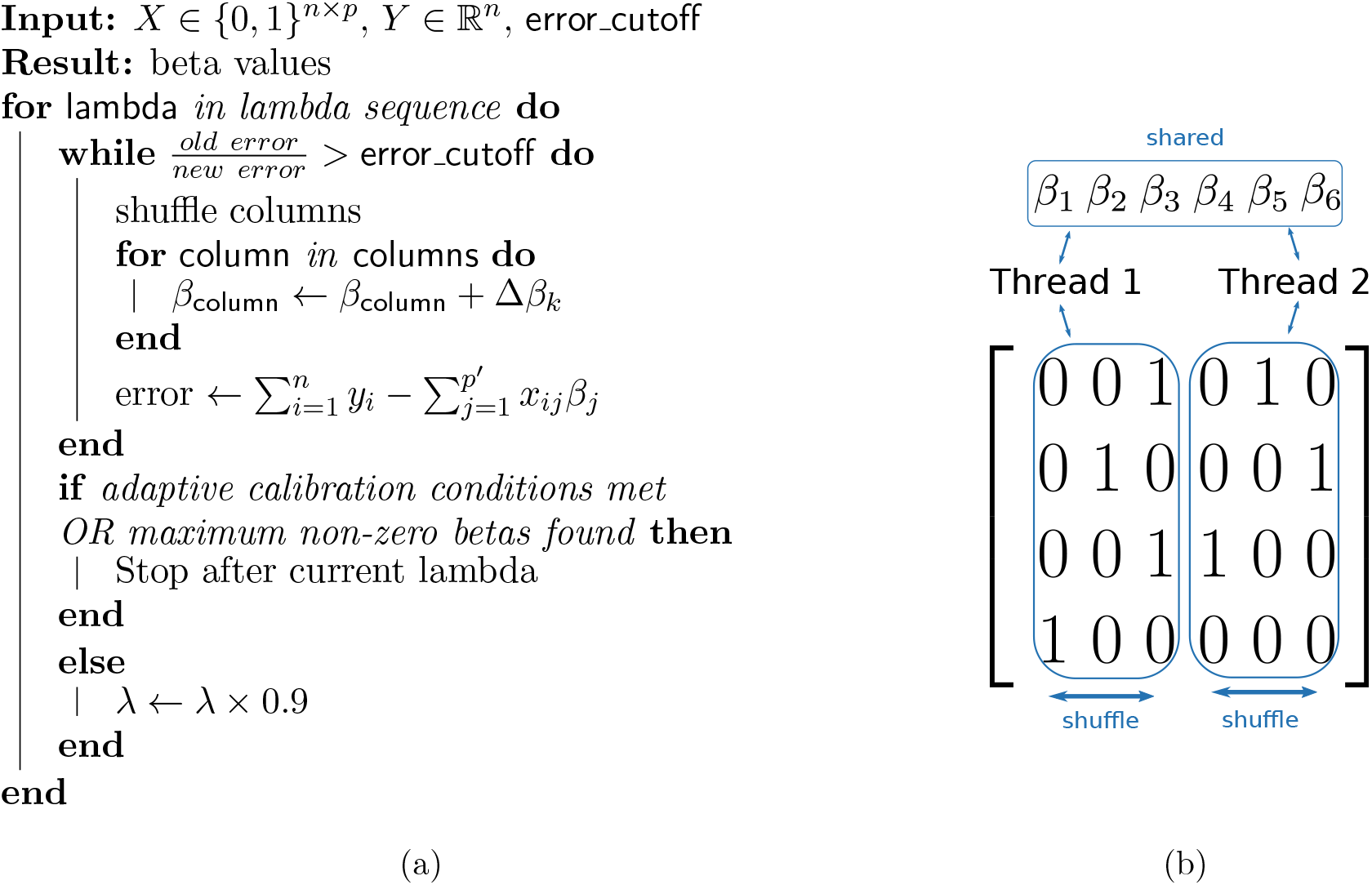
Shuffled Lasso Algorithm (a). Parallel implementation (b).

#### D.5.1. Shuffling Method

For lasso regression on *k* cores, our aim in shuffling the columns is to ensure that columns are not updated at the same time in many iterations. To improve performance when *p* is extremely large, we shuffle the columns in several simultaneous batches, one for each thread.

We therefore implemented a parallel variation of the Fisher and Yates [13] algorithm, as described in [11]. We begin by dividing the *n* columns into 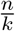 chunks, where *k* is the number of available threads. When iterating through columns in Algorithm 3, these are exactly the chunks that each thread will process. Moving a column from one chunk into another effectively moves it from one thread’s workload into another’s. This is unnecessary and potentially harms cache performance, so we only shuffle within a chunk. As long as the total number of columns is significantly larger than the number of available threads,^7^ which column in a chunk a thread is working on is a random sample from a large list, and the combinations are unlikely to often repeat themselves. This is sufficiently random to avoid the overcorrection effect observed in Fig. 11.

#### D.5.2. Final parallel vs sequential performance

Running this shuffled version (Algorithm 3) on a 16-core NUMA system (two 8-core CPUs with 2-way SMT), we see a reasonable speedup using up to eight cores, with continuing improvements up to 15 threads on the same cores (Fig. 19). Surprisingly, we see a slight drop in performance adding the 16th thread, which is the final available thread on the first NUMA node. This could be because the columns don’t divide as evenly among 16 threads as they do among 15, or simply that there is some noise in the time taken on a busy system. Above 16 threads we don’t see any further improvement, the performance actually worsens. Given that this is the first core of the second CPU, this appears to be the result of slower memory access on the second node, and the overhead of synchronising beta updates across nodes.^8^

**Figure 19.**
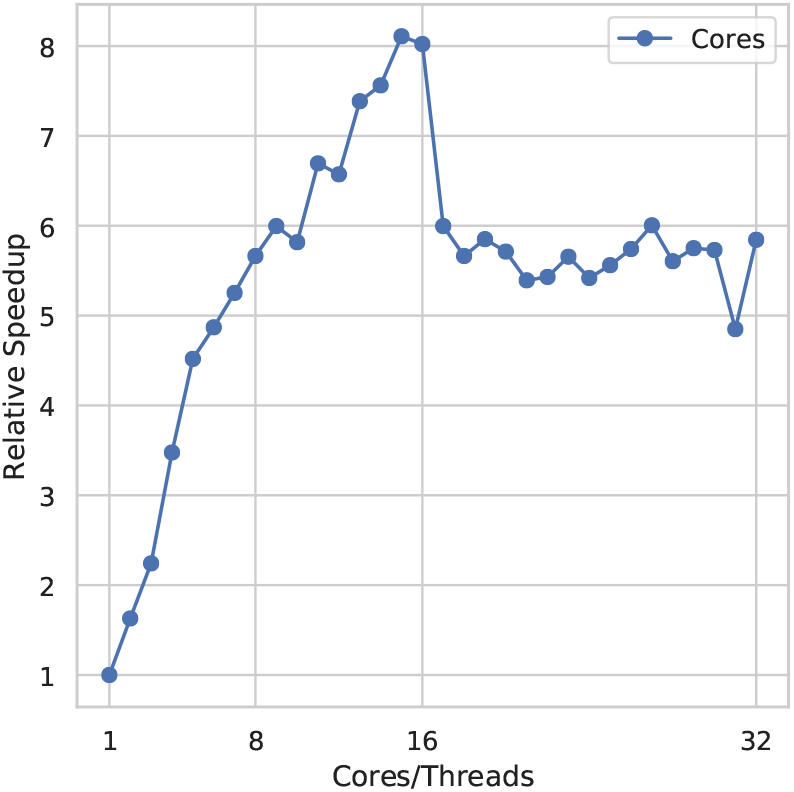
Relative speedup as the number of cores used increases, running on a dual 8 core/16 thread NUMA system. Cores 1-8 are separate cores on node 1, 8-16 are SMT threads on the same cores. Cores 17-24 are separate cores on the second NUMA node, and 15-32 are SMT threads on those cores.

## Footnotes

1 Finding interactions in an siRNA screen of 1,000 genes with ten siRNAs per gene takes several days using ten cores on an Opteron 6276.

2 The compression time is not comparable for all methods. Our Simple-8b implementation compresses columns in parallel, whereas TurboPFor does not. Regression is done in parallel using all cases, using the method described in Appendix D.5.

3 https://github.com/bioDS/Pint/tree/error_compin

4 Assuming that the column updates are done in parallel, and the overcorrection adjustments are done on the main thread.

5 Note that 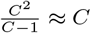 for sufficiently large C

6 Note that it is not the case that one thread is still updating while the others are waiting.

7 And we can assume this true in genome-scale data, with anywhere from thousands to billions of columns

8 Note that steps have been taken to avoid sharing cache lines between CPUs, and each component of the compressed matrix is only accessed by the thread that allocated it.

